# Hair Follicle Epidermal Stem Cells Define a Niche for Tactile Sensation

**DOI:** 10.1101/353490

**Authors:** Chun-Chun Cheng, Ko Tsutsui, Toru Taguchi, Noriko Sanzen, Asako Nakagawa, Kisa Kakiguchi, Shigenobu Yonemura, Chiharu Tanegashima, Sean D. Keeley, Hiroshi Kiyonari, Yasuhide Furuta, Yasuko Tomono, Fiona M. Watt, Hironobu Fujiwara

**Author notes:** Present address: Simmons Comprehensive Cancer Center, UT Southwestern Medical Center, Texas 75390, USA. Present address: Department of Physical Therapy, Niigata University of Health and Welfare, Niigata 950-3198, Japan (Toru Taguchi). Co-first author (Chun-Chun Cheng, Ko Tsutsui, Toru Taguchi).

## Abstract

The heterogeneity and compartmentalization of stem cells is a common principle in many epithelia, and is known to function in epithelial maintenance, but its other physiological roles remain elusive. Here we show transcriptional and anatomical contributions of compartmentalized epidermal stem cells (EpSCs) in tactile sensory unit formation in the hair follicle (HF). EpSCs in the follicle upper-bulge, where mechanosensory lanceolate complexes (LCs) innervate, express a unique set of extracellular matrix (ECM) and neurogenesis-related genes. These EpSCs deposit an ECM protein EGFL6 into the collar matrix, a novel ECM that tightly ensheathes LCs. EGFL6 is required for the proper patterning, touch responses, and αv integrin-enrichment of LCs. By maintaining a quiescent original EpSC niche, the old bulge, EpSCs provide anatomically stable HF-LC interfaces, irrespective of the stage of follicle regeneration cycle. Thus, compartmentalized EpSCs provide a niche linking the HF and the nervous system throughout the hair cycle.

## Introduction

Tissue stem cells (SCs) in many epithelia, including the epidermis, intestine, mammary glands and lungs, often comprise heterogeneous populations with distinct transcriptional and anatomical features (Donati & Watt, 2015; Xin, Greco, & Myung, 2016). Individual SC pools contribute to tissue maintenance through different homeostatic and regenerative properties and give rise to distinct epithelial compartments, while they also hold plasticity to change their identity and function when perturbed. These recent findings question the traditional view of the role of stem cell heterogeneity, which underlies the theory of linear hierarchy (Goodell, Nguyen, & Shroyer, 2015). Although the biological significance of the heterogeneity and compartmentalization of epithelial SCs has primarily been studied in the context of epithelial tissue maintenance and regeneration, its other physiological roles remain unclear.

Hair follicles (HFs) are highly conserved touch sensory organs in the hairy skin that covers most of the mammalian body surface and detects touch signals essential for development and survival (Lumpkin, Marshall, & Nelson, 2010). HFs are innervated by mechanosensory end organs called lanceolate complexes (LCs), which are composed of parallel, longitudinally aligned low-threshold mechanoreceptor (LTMR) axonal endings and terminal Schwann cell (tSC) processes (Zimmerman, Bai, & Ginty, 2014). HFs are also connected to arrector pili muscles (APMs), thus forming a unique regenerating motile sensory organ. These two HF appendages attach to the permanent portion of the HF, known as the bulge, where epidermal stem cells (EpSCs) reside.

Hair follicle EpSCs are heterogeneous in their molecular and functional properties and compartmentalized along the longitudinal axis of the HF (Figure 1A) (Solanas & Benitah, 2013). The mid-bulge region contains CD34^+^ slow-cycling EpSCs that serve as a reservoir of EpSCs and were once thought to be the master SCs at the top of the epidermal hierarchy. A series of recent studies, however, identify several additional pools of different EpSCs around the bulge (e.g. in the isthmus, junctional zone, upper-bulge and hair germ). For example, the upper-bulge is a niche for *Gli1*^+^ EpSCs and they contribute to HF regeneration and wound healing of interfollicular epidermis (Brownell, Guevara, Bai, Loomis, & Joyner, 2011). The importance of EpSC heterogeneity and compartmentalization has almost entirely been explained by their role in regional tissue replenishment to maintain different functional compartments of the epidermis, but its roles apart from epidermal maintenance remain poorly understood. As coordinated epithelial-mesenchymal interactions are essential for HF development, regeneration and functioning, the unique signaling territories and tissue architecture provided by compartmentalized EpSCs probably regulate the interactions between the epidermis and a variety of HF-associated structures, including sensory nerves, APMs and the dermal papilla. Consistent with this idea, it has been reported that mid-bulge EpSCs create a specialized basement membrane containing nephronectin, thereby providing a niche for APM development and anchorage (Fujiwara et al., 2011). Follicle epidermis-derived BDNF is also critical for follicle-nerve interactions (Rutlin et al., 2014).

**Figure 1.**
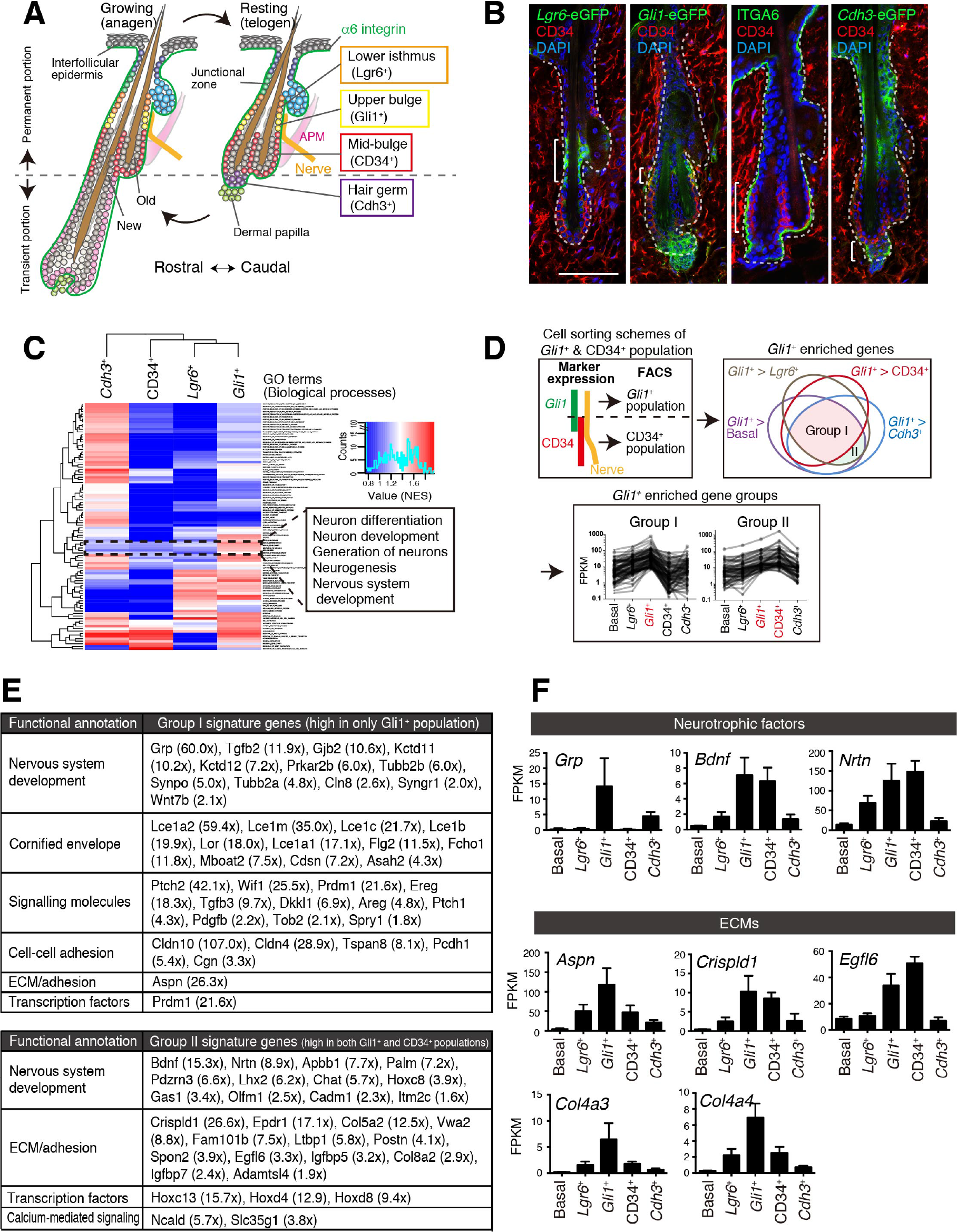
Upper-bulge EpSCs are molecularly specialized for HF-nerve interactions. (**A**) Graphical illustration of EpSC compartments. APM, arrector pili muscle. (**B**) Distinct EpSC compartments in 8-week-old telogen skin were visualized with specific eGFP reporters and cell surface markers. Brackets indicate target cell populations for sorting. (**C**) Z-score heat map representing the normalized enrichment score (NES) of GSEA using the transcriptome data of each EpSC population. (**D**) Scheme of the extraction of genes highly expressed in the *Gli1*^+^ population (Group I) and in both the *Gli1*^+^ and CD34^+^ populations (Group II). (**E**) Lists of the genes highly expressed in *Gli1*^+^ population (Group I) and in both *Gli1*^+^ and CD34^+^ populations (Group II). (**F**) Expression levels of neurotrophic factor and ECM genes highly expressed in upper-bulge SCs. Basal, basal EpSC pool. Data are mean ± SD, n = 3-4.

In this study, we set out to address the idea that upper-bulge EpSCs are molecularly and anatomically specialized for interaction with LCs and thus involved in generating the sense of touch. We demonstrated that upper-bulge EpSCs produce a special extracellular matrix (ECM) environment and unique epidermal tissue architecture that creates a stable HF-LC interface for tactile sensation. Thus, EpSC heterogeneity and compartmentalization appear to be important for both regional epidermal maintenance and patterned epithelial-mesenchymal interactions.

## Results

### Upper-bulge EpSCs are molecularly specialized for HF-nerve interactions

We first examined the global transcriptional features of distinct EpSC populations in the HF. To this end, we established FACS-based cell purification methods using several eGFP reporter mouse lines that label different SC compartments to isolate cellular subpopulations resident in the lower-isthmus (*Lgr6*^+^), upper-bulge (*Gli1*^+^), mid-bulge (CD34^+^), and hair germ (*Cdh3*^+^) as well as unfractionated basal EpSCs (α6 integrin^+^) (Figure 1A and B, Figure 1-figure supplement 1A). The purity of the isolated populations was verified by the expression of compartment-specific genes (Figure 1-figure supplement 1B and C, Materials and methods). We performed RNA-seq on these isolated cell populations. Gene Set Enrichment Analysis (GSEA) indicated that neurogenesis-related Gene Ontology (GO) terms are over-represented in *Gli1*^+^ upper-bulge EpSCs (Figure 1C, Figure 1- table supplement 1). To further identify *Gli1*^+^ compartment-enriched genes, we performed a pairwise transcriptional comparison between the *Gli1*^+^ population and all the other populations and plotted the relationship between *Gli1*^+^-enriched gene sets in a Venn diagram (Figure 1D). Genes categorized in “Group I” were *Gli1*^+^ population-enriched genes. We also extracted genes included in “Group II”, which contains genes highly expressed both in the *Gli1*^+^ population and the CD34^+^ population, since the *Gli1* and CD34 double positive cells were included in the CD34^+^ population in our sorting scheme (Figure 1D). Prominent gene-annotation clusters in both Group I and Group II cells encode proteins involved in nervous system development, including the neurotrophic factors *Grp, Bdnf* and *Nrtn* and the keratitis-ichthyosis-deafness syndrome gene *Gjb2* (Figure 1E and F). Multiple ECM genes are also upregulated in the upper-bulge compartment, including *Aspn, Crispld1, Egfl6*, and the deafness-related ECM genes *Col4a3* and *Col4a4* (Mochizuki et al., 1994) (Figure 1E and F). This global gene expression profiling of compartmentalized EpSCs suggests that upper-bulge EpSCs are specialized both to interact with the nervous system and to express a unique set of ECM genes.

### Upper-bulge EpSCs deposit EGFL6 into the collar matrix

It has been suggested that the ECM plays important roles in mammalian touch end organs, but the molecular identity and functions of this putative ultrastructure remain unknown (Lumpkin et al., 2010; Zimmerman et al., 2014). On examining the tissue localization of 15 upper-bulge ECM proteins, we found that 8 ECM proteins were deposited in the upper-bulge (Figure 2A, Figure 2-table supplement 1). Among them, EGFL6 exhibited the most restricted localization in the upper-bulge of all types of dorsal HFs and showed a unique C-shaped pattern with a gap at the rostral side of the HF (Figure 2B). βIII-tubulin staining showed that skin nerve endings terminate at the EGFL6 deposition sites (Figure 2B). Magnified 3D images revealed the close association of EGFL6 with longitudinal lanceolate parallel LTMR axonal endings of LCs, which are activated by tactile stimuli (Figure 2C) (Bai et al., 2015), and longitudinal processes of nestin-positive non-myelinating tSCs of LCs (Figure 2D). A close anatomical association of follicle EGFL6 with blood vessels has been reported (Xiao et al., 2013), but their direct contact was not observed (Figure 2E). To detect cells expressing *Egfl6* mRNA, we generated *Egfl6-H2b-Egfp* reporter mice. eGFP protein expression was enriched in upper-bulge EpSCs at the caudal side of the HF, but was not detected in dermal cells around the upper-bulge or dorsal root ganglia neurons that innervate to the upper bulge (Figure 2F and G).

**Figure 2.**
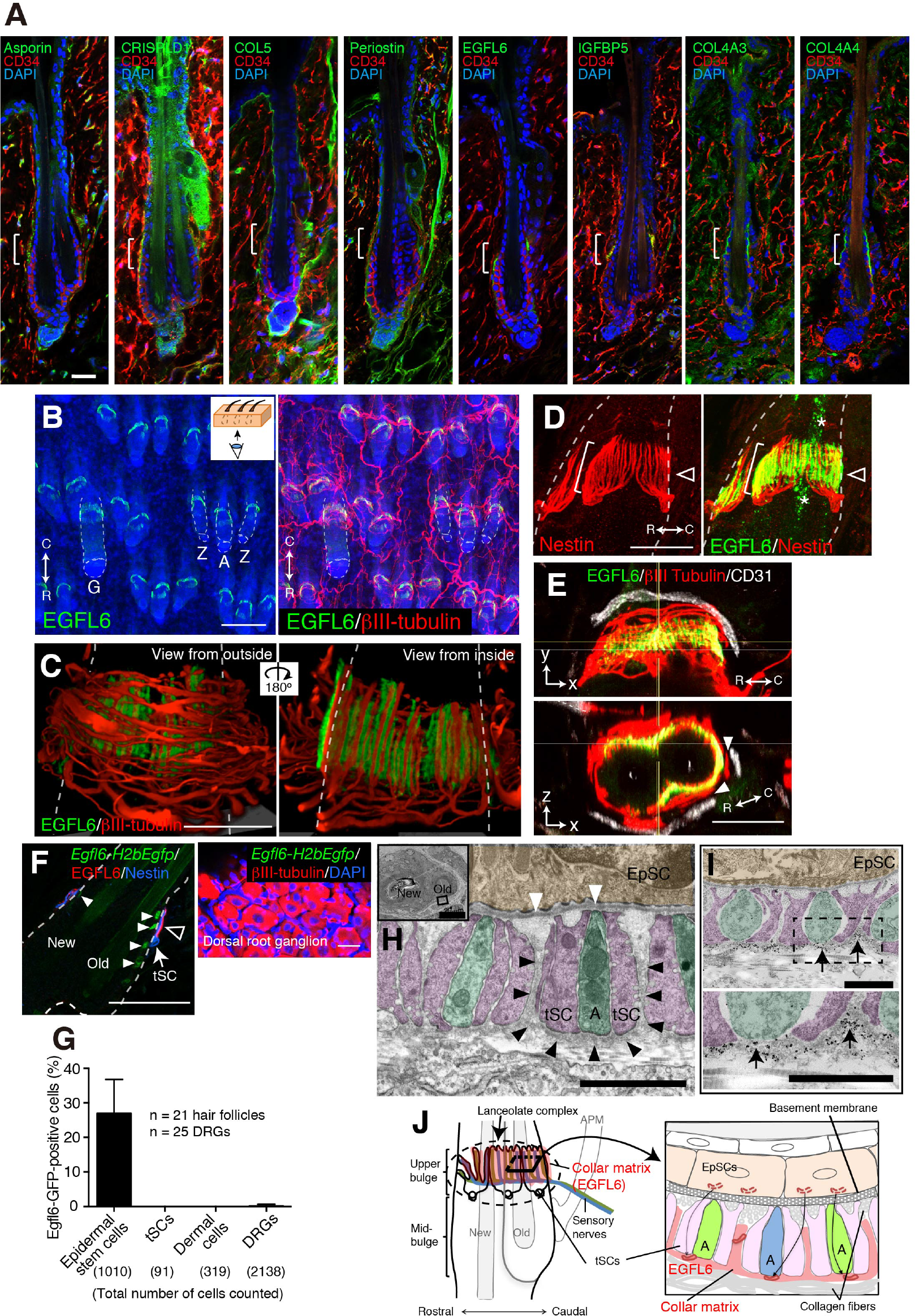
Upper-bulge EpSCs deposit EGFL6 into the collar matrix. (**A**) Immunostaining pattern of upper-bulge-specific ECM in 8-week-old telogen skin. Brackets: upper-bulge. (**B**) EGFL6 was colocalized with βIII-tubulin^+^ nerve endings in the upper-bulge of dorsal guard (‘G’), awl/auchene (‘A’) and zigzag (‘Z’) HFs. (**C**) Magnified images of EGFL6 and βIII-tubulin in telogen skin. (**D**) The protrusions of the tSCs (white brackets) were colocalized with EGFL6 in the upper-bulge of 8-week-old telogen HF (open arrowheads). Asterisks: nonspecific signals. (**E**) CD31^+^ blood vessels did not show direct contact to EGFL6 in 7-week-old telogen HF. Arrowheads: gaps between EGFL6 (green) and blood vessels (white). (**F**) The dorsal skin and dorsal root ganglia (DRG) of an 8-week-old *Egfl6-H2b-Egfp* mouse was stained for eGFP, EGFL6, nestin and βIII-tubulin. eGFP (closed arrowheads) staining was observed in the upper-bulge (open arrowhead) epidermis of old bulge (Old), but not in other cellular components around the upper-bulge. (**G**) Statistical examination of *Egfl6*-H2BeGFP-positive cells around the upper bulge. Data are mean ± SD. (**H**) A transmission electron microscopic image of the upper-bulge of P35 zigzag HF. An electron-dense amorphous ECM structure, the collar matrix (black arrowheads), ensheathed LC endings [A, axon (green); tSC (pink); EpSC (gold); white arrowheads, epidermal basement membrane]. (**I**) The EGFL6-gold particles were localized in the collar matrix (arrows). (**J**) Schematic representation of the localization of EGFL6 in the upper-bulge. Scale bars, 10 μm (A), 20 μm (C-F), 100 μm (B), 2 μm (H), 1 μm (I).

To investigate the ultrastructural localization of EGFL6, we performed electron microscopic analysis of transverse sections of the upper-bulge. As described by Li and Ginty (2014), each LTMR axonal ending was sandwiched by processes of tSCs (Figure 2H). We identified an electron-dense amorphous ECM structure surrounding the LCs (black arrowheads in Figure 2H). We named this novel structure the ‘collar matrix’, as it tightly ensheathes the LCs. tSC processes have two openings, at the follicular basement membrane and at the opposite side of the cell, thus exposing axonal endings and tSC processes to at least two different ECMs, the HF basement membrane and the collar matrix. Although EGFL6 is expressed by the upper-bulge EpSCs, an immunogold-labeled EGFL6 antibody was detected exclusively in the collar matrix (Figure 2I), indicating that upper-bulge EpSCs secrete EGFL6, which is incorporated in the collar matrix that ensheathes LCs (Figure 2J).

### EGFL6 mediates cell adhesion via αv integrins

α8β1 integrin is the only reported cell adhesion receptor of EGFL6 (Osada et al., 2005), but its expression was not detected in LCs (data not shown). To explore other cell adhesion receptors for EGFL6, we purified recombinant EGFL6 protein and performed cell adhesion assays using nine cell lines. Of these, two glial and one skin fibroblast cell lines adhered to EGFL6, whereas other cell types, including keratinocytes, did not (Figure 2-figure supplement 1A and B), suggesting that although EGFL6 is derived from the upper-bulge EpSCs, these SCs are incapable of interacting with EGFL6, rather dermal cell lineages interact and adhere to EGFL6. A point mutation into the Arg-Gly-Asp (RGD) integrin recognition sequence of EGFL6 (RGE mutant) significantly reduced its cell adhesive ability (Figure 2-figure supplement 1C). Cell adhesion to EGFL6 was inhibited by the αv integrin antibody, but not by the β1 integrin antibody (Figure 2-figure supplement 1D). Consistent with these results, αv integrin accumulated in LCs and colocalized with EGFL6 (Figure 2-figure supplement 1E). Deletion of EGFL6 decreased αv integrin and impaired its accumulation (Figure 2-figure supplement 1E and F). Together, these results demonstrate that EGFL6 induces cell adhesion via αv integrins that accumulate in the LCs in an EGFL6-dependent manner.

### EGFL6 is required for the proper patterning and touch responses of LCs

One of the central roles of the specialized ECM around neural structures, such as the perineuronal nets, is to provide adhesive and non-adhesive physical structures to support cells and therefore determine their shape and functions (Mouw, Ou, & Weaver, 2014). Thus, we next examined the effect of deleting EGFL6 on LC structures in zigzag HFs, which are the most common HF type in rodent skin and innervated by δ5- and C-LTMRs (Li et al., 2011). *Egfl6* knockout mice showed misaligned and overlapping structures of axonal endings and tSC processes (Figure 3A and B). Quantitative 3D histomorphometrical analysis of LTMRs and tSCs revealed the significant increase of overlapping points of axons and tSC protrusions in *Egfl6* knockout mice (Figure 3A-C). The numbers and lengths of axonal endings were unchanged (Figure 3-figure supplement 1A and B). In *Egfl6* knockout tissue, the length, but not the width, of tSC processes was reduced (Figure 3-figure supplement 1C and D). *Egfl6* knockout mice did not show defects in the hair cycle (Figure 3-figure supplement 1E and F). We conclude that EGFL6 is required for proper parallel patterning of LCs.

**Figure 3.**
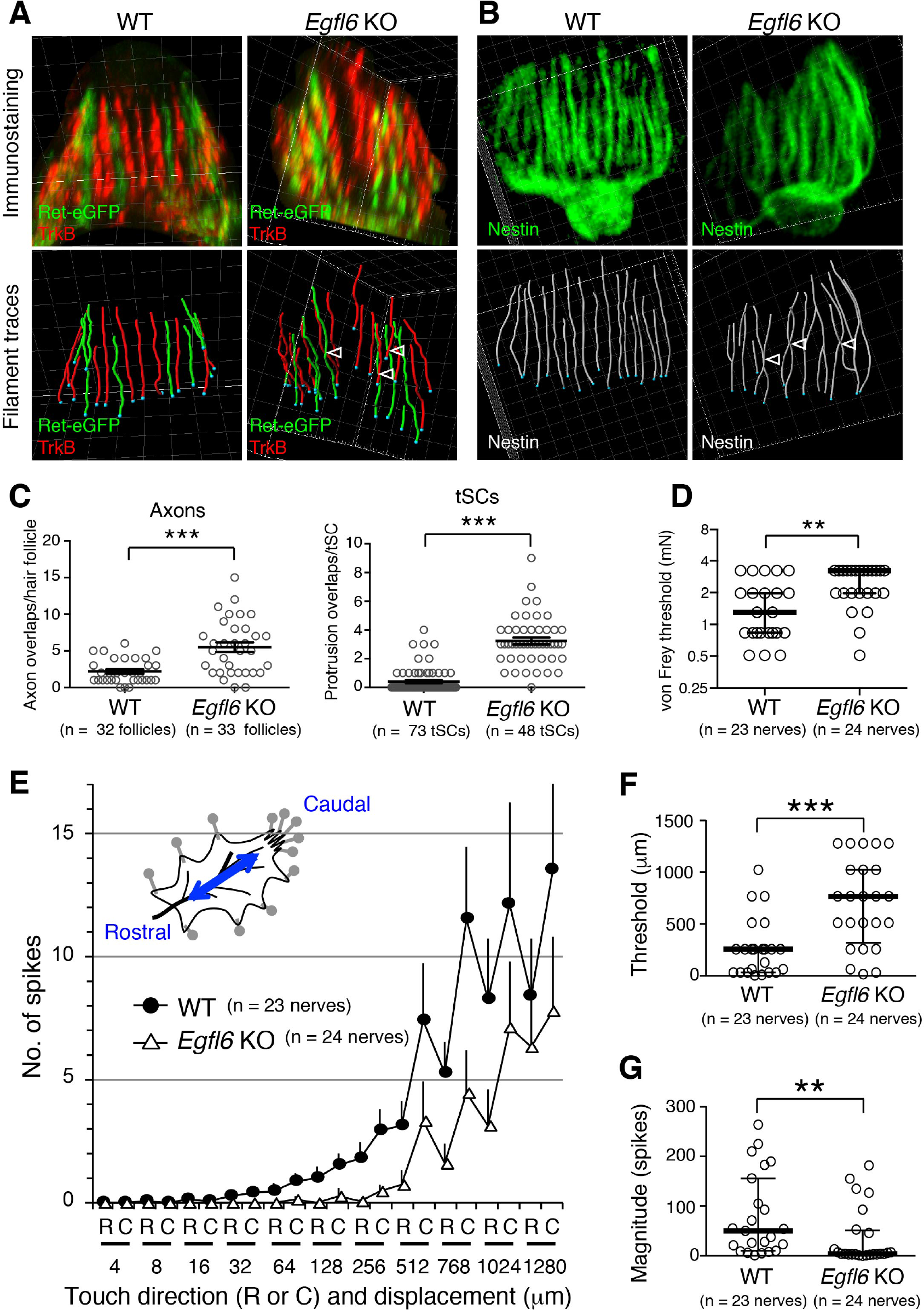
EGFL6 is required for the patterning and touch responses of LCs. (**A** and **B**) 3D reconstituted images and filament tracings of axonal endings (**A**) and tSCs (**B**) in the old bulge of wild-type and *Egfl6* knockout 8-week-old skin. *Ret*-eGFP, a marker for C-LTMRs and TrkB, a marker for Aδ-LTMRs. Open arrowhead: overlapping points of axonal endings and tSC processes. Grid width, 5 μm. (**C**) Overlapping points of axonal endings and tSC processes were counted. Data are mean ± SEM. Two-tailed unpaired *t*-test. (**D**) Mechanical sensitivity of Aδ-LTMRs was analyzed using von Frey hairs in wild-type and *Egfl6* knockout skin-nerve preparations from 10-12 week-old telogen skin. (**E**) Touch response patterns of Aδ-LTMRs in wild-type and *Egfl6* knockout skin-nerve preparations. A ramp-and-hold touch stimulus was applied using a piezo-controlled micromanipulator (see Methods and Figure 3-figure supplement 2A). (**F**) Threshold displacement of Aδ-LTMRs. (**G**) Number of action potentials during the course of the touch stimulus protocol (i.e., magnitude of touch responses) was measured. Data in (D, F, G) are median with interquartile range (IQR) and in (**E**) are mean ± SEM. Statistics: Mann-Whitney U-test (D, F, G).

To determine whether EGFL6 is involved in mechanotransduction, we generated skin-nerve preparations and performed a single-nerve electrophysiological analysis of Aδ-LTMRs, which are the most sensitive HF-associated LTMRs, stimulated by gentle touch and hair deflection (Bai et al., 2015; Rutlin et al., 2014; Zimmermann et al., 2009). Although the electrical responsiveness of Aδ-LTMRs in *Egfl6* knockout skin remained intact (Figure 3-table supplement 1), these receptors showed clear defects in mechanical responses. The median mechanical response threshold of Aδ-LTMRs measured using von Frey hairs was significantly higher in knockout skin (Figure 3D). To more quantitatively analyze the mechanical sensitivity of these fibers, a light touch stimulus was applied using a piezo-controlled micromanipulator (Figure 3-figure supplement 2A). Averaged touch response patterns showed that knockout skin exhibits fewer action potentials than wild-type skin at displacements of >32 μm (Figures 3E, Figure 3-figure supplement 2B and C). The median threshold displacement of Aδ-LTMRs was about three-times higher in the knockout (Figure 3F). Significantly fewer action potentials were detected during the course of the touch stimulus protocol in *Egfl6* knockout skin (Figure 3G). These results indicate that EGFL6 plays an important role in the excitation of Aδ-LTMRs.

### The bulge provides stable epidermal-neuronal interfaces

Unlike most other sensory systems, the HF undergoes dynamic structural changes during its periodic regeneration cycle (Figure 3-figure supplement 1E). It has remained unclear how HFs are able to maintain stable HF-LC interactions over the course of this cycle. To gain insight into this question, we first examined the changes that occur in the anatomical structure of the upper-bulge epidermis during the hair regeneration cycle. We found that at the first telogen (P20), the upper-bulge exhibits a single circular peripheral morphology with one hair shaft, while at the first anagen (P35), the upper-bulge perimeter doubles due to the formation of a new HF rostral to the existing HF (old bulge) (Figure 4A-C). At the second telogen (P49), the upper-bulge perimeter decreased due to a reduction in the new bulge perimeter. Thus, the external configuration of the rostral side of the upper-bulge epidermis changes dynamically during the HF regeneration cycle, while the caudal aspect remains stable.

**Figure 4.**
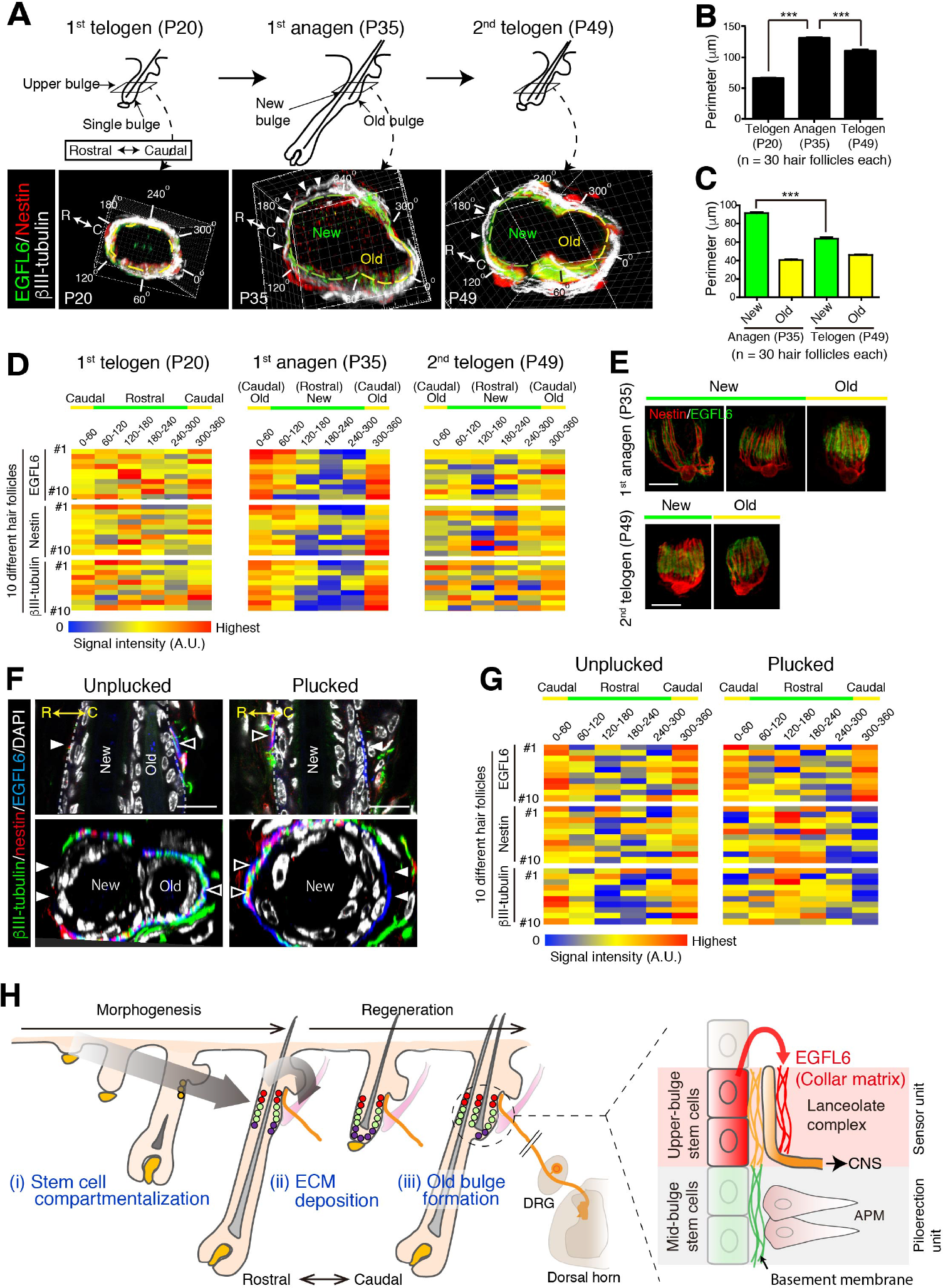
The old bulge provides stable epidermal-neuronal interfaces. (**A**) Structural changes of the upper-bulge during the HF regeneration cycle. Cross-sectional views of the upper-bulge of zigzag HFs are shown. Closed arrowheads indicate a large gap of LCs. (**B**) Perimeter length of the upper-bulge and (**C**) the upper-bulge of new and old bulges. (**D**) Heat map of the relative signal intensity levels of EGFL6, nestin and βIII-tubulin along the upper-bulge perimeter. Geometry of the upper-bulge perimeter is shown above the heat map. (**E**) Morphological differences of EGFL6/tSC complexes in new and old bulges. (**F**) Sagittal and transverse sectional views of the upper-bulge in P35 anagen HFs with the club hair unplucked and plucked. Open and closed arrowheads indicate areas with and without LCs, respectively. (**G**) Heat map of the relative signal intensity levels of EGFL6, nestin and βIII-tubulin along the upper-bulge perimeter in HFs with the club hair unplucked and plucked. (**H**) Schematic summary of the contribution of EpSCs to sensory unit formation. (i) During development, EpSCs for HF-nerve interactions are induced and compartmentalized in the upper-bulge. (ii) Upper-bulge EpSCs provide specialized ECM and neurogenetic environments for LCs. (iii) EpSCs maintain an old bulge at the caudal side of the HF, providing stable HF-nerve interfaces. Scale bars, 10 μm.

We next examined the relationship between the dynamics of HF epidermal morphology and LC structure. We measured the signal intensities of immuno-stained LC components along the upper-bulge, and assigned average signal intensities to the position angle of HFs. At the first telogen, HFs exhibited relatively regular signal intensities along the epidermal perimeter, but at the first anagen the new bulge (rostral side) showed areas of low signal intensity, while these remained unchanged at the old (caudal) bulge (Figure 4A and D). The low-intensity areas at new bulges became narrower in the second telogen. Both Aδ- and C-LTMRs exhibited caudally polarized distribution in the second telogen (Figure 4-figure supplement 1A). Consistent with these signal intensity data, EGFL6 and tSCs in the old bulge maintained regular parallel morphologies throughout the hair cycle, while those in the new bulge exhibited diverse morphologies and wider widths in the first anagen (Figure 4E, Figure 4-figure supplement 1B). Thus, the polarized distributions of LC components toward the caudal side of the HFs are induced as the new bulge structure forms at the rostral side of the HF. The caudal side of the HF, where the old bulge is maintained by quiescent EpSCs, does not change in morphology, suggesting that the old bulge stably preserves LCs irrespective of the stage of hair regeneration cycle.

Finally, we plucked a club hair, a retained hair from the previous hair cycle in the old bulge, to test whether the removal of the sensor probe and subsequent alteration of the old bulge architecture would affect the LC structure. The hairs were painted at the first telogen (P20) and the painted hairs (i.e., club hairs) were plucked when the new hair and bulge formed (Figure 4-figure supplement 1C). Three days after plucking, the old bulges regressed (Figures 4F, Figure 4-figure supplement 1D), the LCs disappeared from the caudal side of the upper-bulge, and ectopic LCs appeared at the rostral side of the upper-bulge (Figure 4F and G). Together these results indicate that the preservation of the old bulge epidermal structure provides a stable epidermal-neuronal interface and induces an LC structure oriented toward the caudal side of the HF. Rutlin et al. (2014) reported that caudally polarized localization of Aδ-LTMRs underlies direction-selective responsiveness of Aδ-LTMRs to hair deflection, suggesting that the unique tissue architecture generated by upper-bulge EpSCs is involved in the directional selectivity of the LCs.

## Discussion

Our findings reveal transcriptional and anatomical roles of compartmentalized EpSCs in HF sensory unit formation (Figure 4H). Our previous study also showed that the mid-bulge EpSCs express many tendon-related genes and induce the differentiation of APMs and their correct anchorage to the bulge via secretion of the basement membrane protein nephronectin (Fujiwara et al., 2011). Deletion of nephronectin moves the muscle near the upper-bulge. The morphology of the LC was altered in the nephronectin mutant (Figure 4-figure supplement 1E-G), likely due to the changes in mechanical or geometrical environments of the upper-bulge by the mis-location of APMs. Therefore, we propose that the heterogeneous EpSCs are compartmentalized not only for efficient epidermal homeostasis and regeneration (Schepeler, Page, & Jensen, 2014), but also for defining patterned niches for specific epidermal-dermal interactions, in part through providing different ECM environments.

The biological significance of forming two bulges has not been well explained. A recent report by Lay, Kume, and Fuchs (2016) suggests that the two-bulge structure has an advantage for life-long HF epidermal maintenance as it increases the number of EpSCs and maintains SC quiescence by holding a keratin 6^+^ inner bulge that expresses factors that inhibit SC activation. Our study now brings an entirely different perspective on the significance of the old bulge formation. We demonstrated that the formation and preservation of the old bulge provides a stable epidermal-neuronal interface and induces a LC structure oriented toward the caudal side of the hair follicle. Caudally polarized expression of BDNF in the bulge epidermis of developing HFs regulates caudally enriched localization of Aδ-LTMRs, which underlies direction-selective responsiveness of Aδ-LTMRs (Rutlin et al., 2014). Our data show that both Aδ-LTMRs and C-LTMRs become caudally polarized after the formation of the two-bulge structure, indicating that the unique tissue architecture provided by EpSCs is another major determinant of the formation, preservation and function of LCs in the HF.

The mechanisms to form and preserve proper innervation in regenerating HF have attracted attention. Botchkarev, Eichmuller, Johansson, and Paus (1997) reported the constant number of longitudinal nerve endings in the LCs in both natural and hair plucking-induced hair cycle. This invariance of nerve ending number, together with the unique epidermal tissue geometry and dynamics revealed in the present study, may underlie stability and topological reorganization of LCs during the hair cycle and in hair plucking-induced old bulge depletion.

We identified a novel electron-dense and amorphous ECM structure, which we named the collar matrix, that ensheathes LCs. The deletion of a collar matrix protein, EGFL6, impaired the structure and sensory function of LCs. The collar matrix has anatomical and functional features in common with the essential ECM in touch end organs of non-vertebrate organisms, such as the mantle in *C. elegans* (Emtage, Gu, Hartwieg, & Chalfie, 2004). Thus, electron-dense amorphous ECMs are likely to be the fundamental ECM regulating sensory end organ formation and function in multicellular organisms.

In conclusion, our findings give a new perspective on the roles played by heterogeneous and compartmentalized EpSC populations in more global aspects of organogenesis, beyond their role in epithelial maintenance and regeneration. Our study also provides insights into how sensory organs take advantage of a stable niche provided by a quiescent EpSC compartment to maintain sensory function within a structurally dynamic tissue environment.

## Author Contributions

C-C.C., K.T. and T.T. contributed equally to the study. C-C.C. and H.F. designed this study. C-C.C., N.S. A.N. and H.F. performed the mouse experiments. H.F., N.S., H.K. and Y.F. generated the *Egfl6-H2b-eGFP* mice. K.T. established the FACS protocols and purified the SC pools. K.T, C.T. and S.D.K. performed the RNA-seq and bioinformatics analysis. T.T. performed the electrophysiological analysis. C-C.C., K.K. and S.Y. performed the electron microscopic analysis. Y.T. provided COL4A3 and COL4A4 antibodies. F.M.W. oversaw the initial stage of this study. C-C.C., K.T., T.T. and H.F. wrote the paper with input from all authors.

## Acknowledgments

We thank Shigehiro Kuraku, Yuichiro Hara, Osamu Nishimura of the RIKEN Phyloinformatics Unit for help in RNA-seq and bioinformatics; Hideki Enomoto for Ret-GFP mice; RIKEN Kobe light microscopy and animal facilities for technical assistance; Shigeo Hayashi and Douglas Sipp for their critical reading of the manuscript. We also thank the members of the Fujiwara laboratory for valuable reagents and discussion. This work was funded by RIKEN intramural funding, JSPS KAKENHI (25122720), Uehara Memorial Foundation, Takeda Science Foundation and Cosmetology Research Foundation (all to H. F.). C-C.C. was a recipient of the RIKEN-NSC Taiwan Fellowship. F.M.W. acknowledges financial support from the Medical Research Council, BBSRC and Wellcome Trust.

## Materials and methods

### Mice

*Egfl6* knockout mice (032277-UCD) were obtained from the Mutant Mouse Resource & Research Centers (MMRRC). Lack of *Egfl6* mRNA and EGFL6 protein were confirmed by *in situ* hybridization and immunostaining. *Egfl6-H2b-Egfp* BAC transgenic mice were generated as described below. *Ret-eGFP* knock-in mice have been described previously (Jain et al., 2006) and kindly provided by H. Enomoto (Kobe University). *Lgr6-GFP-ires-CreERT2* mice were obtained from Jackson Laboratory. *Gli1-eGFP* mice (STOCK Tg(Gli1-EGFP)DM197Gsat/Mmucd) and *Cdh3-eGFP* mice (STOCK Tg(Cdh3-EGFP)BK102Gsat/Mmnc) were obtained from MMRRC. Mouse lines used for transcriptome analysis were backcrossed with C57BL/6N mice more than 4 times. C57BL/6N and BALB/c mice were obtained from Japan SLC Inc. *Npnt* floxed mice were obtained from Jackson Laboratory and crossed with keratin 5-Cre mice from the Jose Jorcano laboratory. All animal experiments were conducted and performed in accordance with approved Institutional Animal Care and Use Committee protocols.

### Generation of the *Egfl6-H2b-Egfp* mouse line

*Egfl6-H2b-Egfp* BAC transgenic mice (Accession No. CDB0518T: http://www2.clst.riken.jp/arg/TG%20mutant%20mice%20list.html) were generated by introducing a BAC carrying a *H2b-Egfp* transgene just before the ATG of the mouse *Egfl6* gene. The BAC clone RP23-124O13, containing the full genomic sequence of *Egfl6*, was obtained from CHORI BACPAC Resources Center. The *H2b-Egfp* fusion gene was introduced just before the first coding ATG of the *Egfl6* gene in the BAC clone using a two-step selection BAC recombineering protocol (Warming, Costantino, Court, Jenkins, & Copeland, 2005). The modified BAC construct was purified with a NucleoBond Xtra Midi kit (Macherey-Nagel) and injected into the pronuclei of fertilized one-cell mouse eggs derived from the breeding of BDF1 and C57BL/6N mice. Potential founder mice were screened by genomic PCR of ear biopsy DNA with following primers: 5’- CCAAGGTCCTGACCAGCGAAG-3’ and 5’- CCTTAGTCACCGCCTTCTTGGAG -3’ (product size: 163 base pairs). These primers were also used for routine genotyping of this mouse line. The transgene-positive founder mice were bred with wild-type C57BL/6N mice and some of the offspring were used for immunostaining of eGFP to confirm the expression and nuclear localization of the H2B-eGFP protein. We have established 16 nuclear-GFP-positive transgenic mouse lines and confirmed the GFP expression in the upper bulge in 13 mouse lines.

### Cell lines

Cell lines used in this study were 293F cells (Thermo Fisher), human primary keratinocytes (Watt lab), human embryonic kidney epithelial cell line HEK293 (RIKEN Bioresource Center), human glioblastoma cell line T98G (JCRB Cell Bank), Human glioma cell line KG-1-C (JCRB Cell Bank), human skin fibroblast cell line SF-TY (JCRB Cell Bank), human embryonic fibroblast cell line NTI-4 (JCRB Cell Bank), human sarcoma cell line HT-1080 (JCRB Cell Bank), human myelogenous leukemia cell line K562 (JCRB Cell Bank) and human myelogenous leukemia cell line transfected with human integrin α8 cDNA K562-a8 (Gift from L. Reichardt).

Human primary keratinocytes were cultured on irradiated J2 feeder cells with DMEM/Ham’s F12 medium supplemented with 10% FBS, 0.5 μg/ml hydrocortisone (Sigma), 0.1 nM cholera toxin (Sigma), 10 μg/ml EGF (Peprotech), 2 mM GlutaMax (Invitrogen) and 5 μg/ml Insulin (Sigma) at 37 °C under 5% CO_2_. Other cell lines were cultured according to the cell culture instructions from each source cell bank or company.

### Antibodies

Antibodies used in this study were listed in Table 1.

**Table 1.** Antibodies used in this study.

| ANTIBODY | SOURCE | IDENTIFIER |
| --- | --- | --- |
| Rabbit anti-mouse EGFL6 | Fujiwara Lab | This paper |
| Rabbit anti-mouse nephronectin | Fujiwara Lab | This paper |
| Mouse anti-keratin 14 | abcam | LL002 (ab77684) |
| Mouse anti-keratin 15 | Santa Cruz | LHK15 (sc-47697) |
| Mouse anti-βIII-tubulin | abcam | 2G10 (ab78078) |
| Chicken anti-nestin | Aves Labs | NES0407 |
| Rat anti-nestin | Millipore | rat-401 (MAB353) |
| Chicken anti-GFP | abcam | ab13970 |
| Rabbit anti-GFP | MBL | 598 |
| Mouse anti-laminin γ1 | Millipore | MAB1914 |
| Rat anti-integrin αv | abcam | RMV-7 |
| Goat anti-TrkB | R&D Systems | AF1494 |
| Rat anti-CD45-PE-Cy7 | eBioscience | 30-F11 |
| Rat anti-TER-119-PE-Cy7 | eBioscience | TER119 |
| Rat anti-CD31-PE-Cy7 | eBioscience | 390 |
| Rat anti-Sca-1-PerCP-Cy5.5 | eBioscience | D7 |
| Rat anti-CD34-eFluor660 | eBioscience | RAM34 |
| Rat anti-CD49f-PE | eBioscience | GoH3 |
| Mouse anti-integrin α2 | DSHB | P1E6 |
| Mouse anti-integrin α3 | DSHB | P1B5 |
| Mouse anti-integrin α5 | DSHB | P1D6 |
| Rat anti-integrin α6 | BioLegend | GoH3 (313614) |
| Mouse anti-integrin αv | ATCC | L230 |
| Mouse anti-integrin β1 | DSHB | P5D2 |

### Generation of antibodies

Rabbit antiserum to mouse EGFL6 was generated by immunizing rabbits with a Myc-His-tagged Mucin-MAM domain of recombinant mouse EGFL6 (S257-G550). The antigen was expressed in the 293F mammalian expression system (Invitrogen) and purified from the supernatant of the transfected cells with a Ni-NTA column. The antibody in the antiserum was affinity-purified with an antigen conjugated CNBr-activated Sepharose 4B. The specificity of the antibody to mouse EGFL6 was confirmed by the absence of the antibody immunoreactivity to tissue samples from *Egfl6* knockout mice. Rabbit antiserum to mouse nephronectin was generated by the same method described above. Recombinant flag-tagged full-length mouse nephronectin was expressed in 293F cells and purified with a FLAG-M2 immuno-affinity column (Sigma). The antibody in the antiserum was affinity-purified with a CNBr-activated Sepharose 4B conjugated with purified His-tagged mouse full-length nephronectin. The specificity of the antibody to mouse nephronectin was confirmed by the absence of the antibody immunoreactivity to tissue samples from *Npnt* knockout mice.

### FACS

Mouse adult dorsal telogen keratinocytes were isolated and stained for cell surface markers as described previously (Fujiwara et al., 2011) with some modifications. We utilized *Lgr6-GFP-ires-CreERT2, Gli1-eGFP* and *Cdh3-eGFP* mice to fluorescently visualize the epidermal stem cells in the lower isthmus (*Lgr6*^+^), upper bulge (*Gli1*^+^), and hair germ (*Cdh3*^+^) with eGFP. Dissected dorsal skin of the 8-week-old female mice was treated with 0.25% trypsin solution (Nakalai tesque) at 37°C for 1 hr. The epidermal tissue was scraped off from the dermal tissue with a scalpel. This epidermis separation protocol leaves hair germ cells in the dermal tissue. Effectiveness of this tissue separation was verified by the following qRT-PCR, RNA-seq and immunohistochemical analyses with compartment-specific markers as described below. For the sorting of lower isthmus (*Lgr6*-eGFP^+^), upper-bulge (*Gli1*-eGFP^+^) and mid-bulge (CD34^+^) epidermal stem cells, the separated epidermis was minced with scalpels and mixed with repeated pipetting to make a single cell suspension. To deplete haematopoietic and endothelial cells (lineage-positive cells; Lin^+^), the cell suspension was stained with PE-Cy7-conjugated antibodies for CD45, TER-119 and CD31. To sort the target cells, the cell suspension was also stained with Sca-1-PerCP-Cy5.5, CD34-eFluor660, CD49f (integrin α6)-PE. Sca-1 was used to remove epidermal basal cells of the interfollucular epidermis and follicle infundibulum (Jensen et al., 2008). Cells were sorted with a FACSAria II according to the expression of cell surface markers, after gating out dead and Lin^+^ cells (Figure 1-figure supplement 1A).

Hair germ epidermal cells (*Cdh3*-eGFP^+^) were sorted from the remaining dermal tissue since the most hair germ epidermal cells were retained in the dermal tissue after the separation of the epidermis. The dermal tissue was minced with scalpels and incubated with 2 mg/ml of collagenase type I at 37°C for 2 hr with gentle mixing. Single cell suspension was obtained by repeated pipetting. The cell suspension was stained with the same antibodies used above and subjected to the sorting procedure.

The purity of sorted cells was confirmed by qRT-PCR with compartment-specific genes: *Lrig1* (junctional zone and lower-isthmus), *Lgr6* (lower-isthmus and upper bulge), *Gli1* (upper-bulge and hair germ), *Bdnf* (upper-bulge), *Cd34* (mid-bulge), *Cdh3* (hair germ) (Figure 1-figure supplement 1B). While the expression of *Gli1*-eGFP was detected in both upper-bulge and hair germ regions (Figure 1B), our cell isolation and sorting methods gave highly pure upper-bulge cells and hair germ cells, respectively. For example, a marker for upper-bulge cells, *Bdnf*, was highly enriched in *Gli1*+ upper bulge cells, but not in *Cdh3*+ hair germ cells (Figure 1-figure supplement 1B). On the other hand, the expression of *Cdh3*, a marker for hair germ cells, was high in the *Cdh3*+ hair germ population, but was very low in the *Gli1*+ upper-bulge population (Figure 1-figure supplement 1B). We identified *Spon1* as a gene exclusively expressed in the hair germ from our gene expression profiling of isolated stem cell populations and confirmed that it was not detected in the *Gli1*+ upper-bulge population (Figure 1-figure supplement 1C). Subsequent protein tissue-localization analysis of compartment-specific genes, which were identified in our gene expression profiling, further confirmed the validity and purity of sorted cells. For example, gene products of upper-bulge-enriched ECM genes, including *Aspn, Crispld1, Egfl6, Col4a3* and *Col4a4*, were deposited in the upper-bulge (Figure 2A), while SPON1 protein was localized in the hair germ region, but not in the upper-bulge. Collectively, our data demonstrate that the purity of each isolated stem cell population was very high.

### qRT-PCR

Total RNA was extracted from the sorted cells using an RNeasy micro kit (Qiagen). qRT-PCR was performed to determine target cell populations using Power SYBR Green PCR Master Mix (Life Technologies) with specific primer sets (Table 2) on 7900HT real-time PCR system (Applied Biosystems).

**Table 2.**
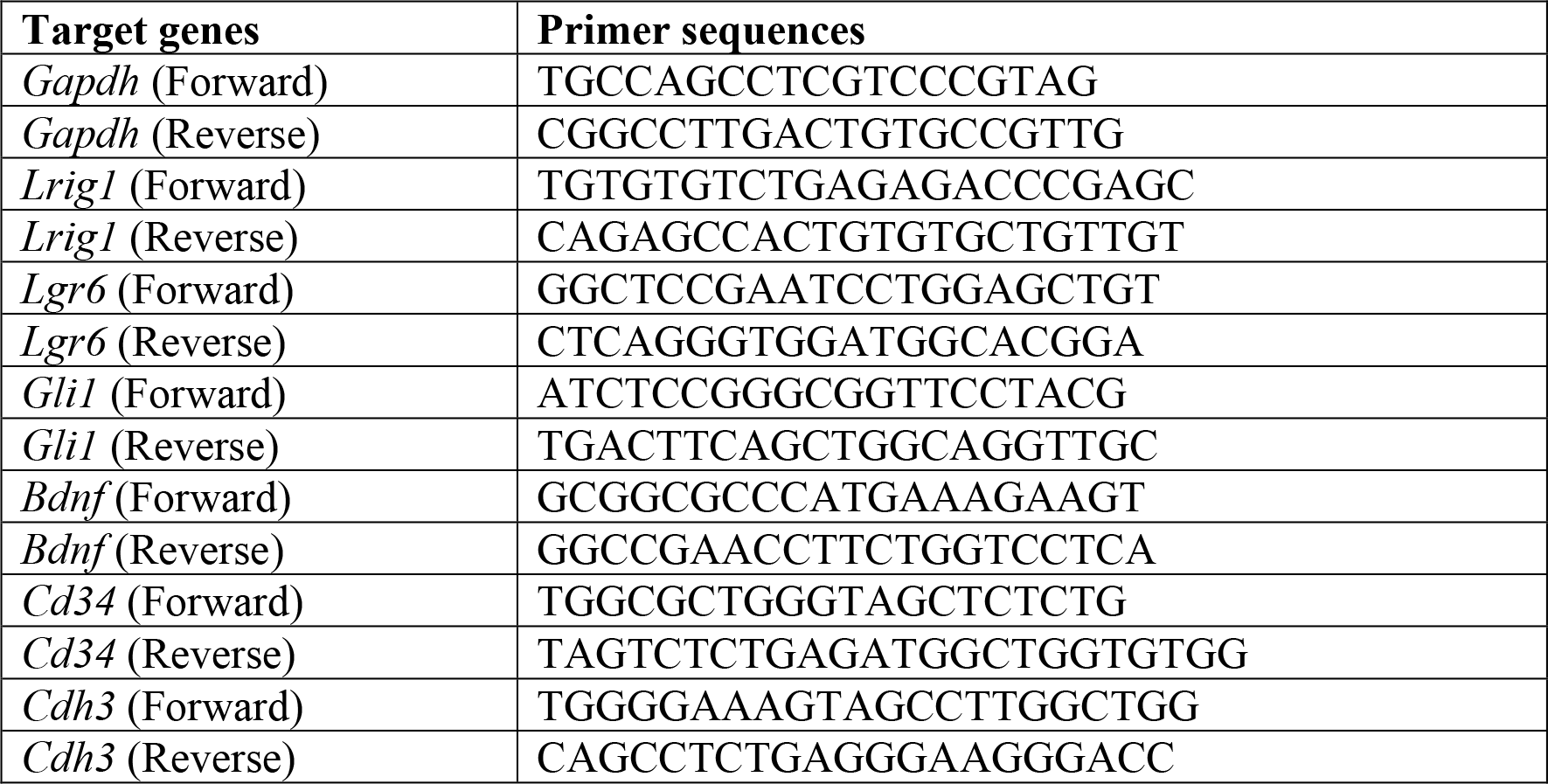
Primer sequences for qRT-PCR.

### RNA library preparation and sequencing

For each library, isolated total RNA samples were quantified with the Qubit RNA HS Assay Kit on a Qubit Fluorometer (Thermo Fisher Scientific), and their qualities were analyzed with the RNA 6000 Pico Kit on a 2100 Bioanalyzer (Agilent Technologies). Qubit measurements were used to ensure that each library was prepared with 10 ng of total RNA. Library preparation was processed following the TruSeq Stranded mRNA Sample Prep Kit (Illumina) protocol till adapter ligation, except that the duration of the initial RNA fragmentation was shortened to 7 minutes. The adapter-ligated cDNA was amplified with 14 PCR cycles. The prepared libraries were sequenced using the Rapid Run mode with 80 cycles on the HiSeq1500 (Illumina), operated by HiSeq Control Software v2.2.58 using HiSeq SR Rapid Cluster Kit v2 (Illumina) and HiSeq Rapid SBS Kit v2 (Illumina). Base calling was performed with RTA v1.18.64 and the fastq files were generated with bcl2fastq v1.8.4 (Illumina).

### Mapping and expression quantification

Qualities of the RNA-seq reads were evaluated with FastQC v0.11.3. The program fastq_quality_filter, part of the FASTX Toolkit v0.0.14, was used to trim bases with a quality value below 30. Additionally, if more than 20% of the bases of a read were removed by this trimming procedure, then the whole read was discarded. Once this quality trim was completed, Trim_galore v0.3.3 was executed to remove adapter sequences from the 3’-ends of reads, and to discard reads that were shorter than 50nt. The processed reads were mapped to the mm10 mouse genome assembly, obtained from iGenomes (http://support.illumina.com/sequencing/sequencing_software/igenome.html), using the splice-aware aligner TopHat v2.0.14 with default parameter settings. Reads mapping to ribosomal DNA accounted for ~1% of the total number of reads in each library, and were removed. Details for each library concerning sequencing and read statistics, including the total amount that uniquely mapped to the genome, are displayed in Table 3.

**Table 3.** RNA-seq read counts and mapping statistics.

|  | Raw<br>Read Count | Trimmed<br>Read Count | rDNA<br>Read Count | Multi-Mapped<br>Read Count | Uniquely Mapped<br>Read Count |
| --- | --- | --- | --- | --- | --- |
| Mouse_Basal_<br>Cells_Rep1 | 11,757,494 | 11,278,061 | 148,700 | 170,077 | 10,456,332 |
| Mouse_Basal_<br>Cells_Rep2 | 11,245,045 | 10,668,752 | 73,729 | 134,777 | 10,051,376 |
| Mouse_Basal_<br>Cells_Rep3 | 11,313,239 | 10,747,103 | 108,596 | 146,602 | 10,089,518 |
| Mouse_Bulge_<br>StemCells_Rep1 | 11,735,034 | 11,181,377 | 64,614 | 183,557 | 10,520,864 |
| Mouse_Bulge_<br>StemCells_Rep2 | 11,917,049 | 11,515,851 | 54,154 | 175,176 | 10,889,932 |
| Mouse_Bulge_<br>StemCells_Rep3 | 13,417,100 | 12,895,113 | 88,655 | 218,495 | 12,090,872 |
| Mouse_Cdh3+_<br>Cells_Rep1 | 10,875,593 | 10,409,342 | 68,193 | 177,407 | 9,574,901 |
| Mouse_Cdh3+_<br>Cells_Rep2 | 11,565,616 | 10,993,093 | 65,001 | 185,981 | 10,079,620 |
| Mouse_Cdh3+_<br>Cells_Rep3 | 11,405,265 | 10,746,768 | 76,143 | 206,384 | 9,790,096 |
| Mouse_Cdh3+_<br>Cells_Rep4 | 11,759,881 | 11,186,543 | 85,902 | 191,154 | 10,274,371 |
| Mouse_Gli1+_<br>Cells_Rep1 | 12,063,488 | 11,478,537 | 57,630 | 178,646 | 10,751,521 |
| Mouse_Gli1+_<br>Cells_Rep2 | 19,762,234 | 18,804,713 | 178,685 | 303,848 | 17,446,776 |
| Mouse_Gli1+_<br>Cells_Rep3 | 19,543,117 | 18,568,695 | 122,017 | 287,214 | 17,378,904 |
| Mouse_Lgr6+_<br>Cells_Rep1 | 20,044,455 | 19,058,807 | 121,507 | 307,804 | 17,798,594 |
| Mouse_Lgr6+_<br>Cells_Rep2 | 19,756,354 | 18,736,071 | 197,458 | 298,652 | 17,386,391 |
| Mouse_Lgr6+_<br>Cells_Rep3 | 19,639,299 | 18,680,969 | 217,502 | 298,146 | 17,210,635 |

Gene expression quantification was performed using the Cuffdiff program in the Cufflinks package v2.2.1, with the frag-bias-correction and multi-read-correction options enabled. Cuffdiff used the mm10 gene model obtained from iGenomes to estimate the number of fragments that originated from individual genes. For each gene, in addition to this produced raw count data, Cuffdiff also calculated normalized expression values, which are referred to as fragments per kilobase of transcript per million mapped reads (FPKM).

### GO and clustering analysis

FPKM values from RNA-seq data were analysed by Gene Ontology (GO) term enrichment analysis. Genes with low expression (FPKM < 5) were filtered out and considered not expressed. Then the remaining FPKM values were log2-transformed for further analysis. To characterize global gene expression profiles of each epidermal stem cell population, gene set enrichment analysis (GSEA) was performed using GSEA software (2.2.2 from Broad Institute) with GO biological processes as a gene set collection (c5.bp.v5.1.symbols.gmt). In this analysis, pairwise differences between basal epidermal stem cells and each epidermal stem cell population were examined. The normalized enrichment score (NES) and False Discovery Rate (FDR) were calculated for each gene set and compared among cell populations using a bioconductor R with heatmap.2 gplots package.

### Immunohistochemistry and imaging

Whole-mount immunostaining and horizontal imaging of mouse dorsal skin were performed as described previously (Fujiwara et al., 2011). Vertical whole-mount imaging of dorsal and whisker skin was performed as described below. Mouse skin tissues were dissected and fixed with 4% paraformaldehyde/PBS for 1 hr at 4°C. Fixed tissues were washed and embedded in OCT compound and frozen on liquid nitrogen. Vertical skin sections (150 μm thick) were made using a cryostat (Leica) and washed with PBS to remove remnant OCT compound. Skin sections were blocked with a blocking buffer (0.5% skim milk/0.25% fish skin gelatin/0.5% Triton X-100/PBS) for 1 hr at 4°C and then incubated with primary antibodies diluted in blocking buffer overnight at 4°C. Skin samples were washed with 0.2% Tween 20/PBS for 4 hr and then incubated with DAPI and secondary antibodies diluted in blocking buffer overnight at 4°C. Finally, skin samples were washed with 0.2% Tween20/PBS for 4 hr at 4°C and mounted with BABB clearing solution. Images were acquired using a Leica TSC SP8. Z stack maximum projection images and three dimensional reconstructed images of skin whole-mount preparation were produced using Imaris 4D rendering software (Bitplane) and Volocity 3D imaging software (Perkin Elmer).

### Transmission electron microscopy

Dissected skin tissues were immediately immersed in 2% fresh formaldehyde and 2.5% glutaraldehyde in 0.1M sodium cacodylate buffer (pH 7.4), sliced into 0.5-1 mm-thin sections with a scalpel and fixed for 2 hr at room temperature. After washing three times with 0.1M cacodylate buffer (pH 7.4) for 5 min, tissues were post-fixed with ice-cold 1% OsO4 in the same buffer for 2 hr. Samples were rinsed with distilled water, stained with 0.5% aqueous uranyl acetate for 2 hr or overnight at room temperature, then dehydrated with ethanol and propylene oxide, and embedded in Poly/Bed 812 (Polyscience). Ultra-thin sections were cut, doubly-stained with uranyl acetate and Reynold’s lead citrate, and viewed with a JEM 1010 or JEM 1400 plus transmission electron microscope (JEOL) at an accelerating voltage of 100 kV.

### Immunoelectron microscopy

Skin samples were dissected, fixed and stained according to the whole-mount immunostaining method. In-house rabbit EGFL6 antibody was used as the primary antibody and Alexa Fluor 488 FluoroNanogold-anti rabbit IgG (Nanoprobes) (1:200 dilution) was used for the secondary antibody. After confirming the localization of Alexa Fluor 488 labelled EGFL6 under a fluorescent microscope, the samples were fixed for 1 hr in 1% glutaraldehyde in PBS. GoldEnhance EM (Nanoprobes) was used in accordance with the manufacturer’s protocol to enlarge the size of gold particles. Samples were then embedded in resin (poly/bed 812; Polyscience). Ultrathin sections were stained with uranyl acetate and lead citrate before observation with an electron microscope (JEM 1400 plus; JEOL).

### Quantification of axon and tSC processes

Neural filaments and tSC protrusions were visualized, measured and analysed using the Imaris software program FilamentTracer (Bitplane). To detect Aδ- and C-LTMRs and tSCs in *Egfl6* knockout mice, we crossed *Egfl6* knockout mice with *Ret-eGFP* knock-in mice (Jain et al., 2006) that express eGFP in C-LTMRs (Li et al., 2011). Skin whole-mounts were immunostained for TrkB, eGFP and nestin to visualize Aδ- and C-LTMRs and tSCs, respectively. To examine the structure of lanceolate complexes, we analysed old bulge regions, but not new bulge regions, since the new bulge regions tend to show a small amount of EGFL6 and an irregular shape of lanceolate complexes. Axon endings and tSC processes were automatically detected in three dimensionally reconstructed immunohistochemical images using FilamentTracer. Traced axonal endings and tSC protrusions were visualized as fixed-diameter cylinders in green (*Ret*-eGFP), red (TrkB) and white (nestin). Length, number and width of axon endings and tSC processes were automatically measured by the software. Overlapped filament points were detected three-dimensionally using the cylinder display mode.

For the measurement of the upper-bulge perimeter, immunostained whole-mount z-stack images were reconstituted to 3D images with Imaris 7.2.1 software. An Imaris Oblique Slicer function was used to make a plane cutting through the region of tSC processes and this plane was defined as a measurement layer. The perimeter of the hair follicle was traced and the length of the trace was automatically measured by the software.

### Quantification of the expressions of *Egfl6*-H2BeGFP and αv integrins

GFP expression in *Egfl6*-H2BeGFP reporter mice were used to examine the expression of *Egfl6* transcripts. Adult telogen dorsal skin of *Egfl6*-H2BeGFP mice were immunostained for GFP, keratin 14 (marker for basal keratinocytes) and nestin (marker for tSCs), with DAPI nuclear counter staining, as described above. Vertical 3D images of zigzag hair follicles were obtained. The number of total and GFP-positive basal epidermal cells, terminal Schwann cells and other dermal cells at the upper-bulge were counted with reference to GFP, keratin-14, nestin and DAPI staining using Imaris software. Twenty-one zigzag hair follicles from three mice were used. The total number of cells counted were 1, 010 basal epidermal stem cells, 91 terminal Schwann cells and 319 other dermal cells.

To investigate the expression of *Egfl6* in sensory nerves of lanceolate complexes, we examined the GFP expression in the adult thoracic dorsal root ganglia (DRGs) of *Egfl6*-H2BeGFP mice since the nuclei of skin sensory nerves are located in the DRGs. DRGs were isolated and fixed with 4% PFA for 1 hr at 4°C. Whole DRG tissues were immunostained for bIII tubulin (marker for DRG neurons) and GFP, with DAPI counter stain. Single sectional plane images were obtained as described above. Twenty-five DRGs from three mice were examined and 2,138 DRG neurons were quantified for their nuclear GFP expression.

To quantify the expression level of αv integrins, we stained telogen adult dorsal skin of wild-type and *Egfl6* knockout mice for EGFL6, αv integrin and nestin, with DAPI counter stain. Signal intensity of αv integrins at the outer surface of the lanceolate complexes was measured using Fiji ImageJ 1.0. Thirteen hair follicles from three wild-type mice and 16 hair follicles from three *Egfl6* knockout mice were used for quantification.

### *Ex vivo* skin-nerve preparations

Receptive properties of cutaneous Aδ-LTMRs were studied using mouse hindlimb skin-saphenous nerve preparations *ex vivo* (Zimmermann et al., 2009). Ten- to twelve-week-old male mice, which were in the resting hair growth phase, were euthanized by inhalation of CO_2_ gas and the preparations were quickly isolated from the left hindlimb. The excised skin was oriented with the dermis side up in the test chamber and affixed with pins. The preparation was maintained at 32 ± 0.3°C (pH 7.4) during the experiment under superfusion with modified Krebs-Henseleit solution (Krebs solution), which contained 110.9 mM NaCl, 4.7 mM KCl, 2.5 mM CaCl_2_, 1.2 mM MgSO_4_, 1.2 mM KH_2_PO_4_, 25 mM NaHCO_3_ and 20 mM glucose. The perfusate was continuously bubbled and equilibrated with a gas mixture of 95% O_2_ and 5% CO_2_. The hindlimb skin of mouse is thinner than dorsal skin.

### Recordings of Aδ-LTMRs

Single Aδ-LTMR was searched and identified when it fulfilled the following criteria: 1) fibers with conduction velocity between 2-10 m/s, 2) fibers responding well to innocuous touch stimulus applied by a blunt glass rod compared to noxious vertical indentation of the skin (compression), 3) fibers without mechanical stimulus intensity-dependent increase in the firing rate, 4) fibers exhibiting a brisk, rapidly adapting discharge at the onset of a supramaximal constant force stimulus (Koltzenburg, Stucky, & Lewin, 1997), 5) fibers not responding to noxious heat and cold stimuli (Figure 3-figure supplement 2) (Zimmermann et al., 2009), and 6) fibers showing a stronger excitation typically in the ramp phase of mechanical stimulation when the stimulus probe was horizontally moving compared to the hold phase, as shown in Figure 3-figure supplement 2. The fibers identified by these criteria in wild type dermis-up skin-nerve preparation exhibited von Frey Hair threshold (vFH) values of ≤0.51-3.23 mN (median: 1.3 mN, IQR: 0.8-2.0 mN) (Figure 3D), which fell into the range of previously reported vFH threshold values of Aδ-LTMRs (or D-hair) with the same dermis-up method (<1-5.7 mN), but not that of high-threshold Aδ-mechano nociceptors (5.7-128 mN) (Zimmermann et al., 2009).

When an Aδ-LTMR was identified, the receptive field (RF) was stimulated with electronic pulses via a bipolar stimulating electrode (frequency of 0.5 Hz, pulse duration of 100 μs and stimulus intensity of <50 V) to measure the conduction velocity of a fiber. The conduction velocity was calculated from the distance and conduction latency of a spike induced by electrical stimulation of the RF. Spontaneous activity of Aδ-LTMRs was analysed for 20 s just before a series of touch mechanical stimulation. Distribution (size and location) of the RFs was mapped on a standardized chart. The size of the RF was measured by calculating the number of pixels in the RF that was drawn on a chart with Image J software. All of the data were stored in a computer via an A/D converter (Power Lab/16s, ADInstruments) with a sampling frequency of 20 kHz. Action potentials were analysed on a computer with the DAPSYS data acquisition system (http://www.dapsys.net). Quantitative mechanical, cold and heat stimuli were then applied to the identified RF in the following order: 1) ramp-and-hold touch mechanical stimulation, 2) cooled from 32°C to 8°C and 3) heated from 32°C to 50°C.

### Mechanical stimulation

At first, the mechanical sensitivity of Aδ-LTMRs was semi-quantitatively analysed using a series of self-made von Frey hairs (VFHs: 0.5-17.6 mN, 0.5 mm in diameter). The strength of the weakest filament that caused a mechanical response was taken as the threshold.

For quantitative analysis of the mechanical sensitivity of a fiber, a light touch stimulus was applied using a piezo-controlled micromanipulator (Nanomoter MM3A, Kleindiek Nanotechnik, Reutlingen, Germany). The stimulator had a glass probe with a spherical tip (diameter: 0.5 mm). After placing the probe tip to the most sensitive point on the identified RF, a series of ramp-and-hold touch mechanical stimuli were applied by driving the stimulator with a pre-programed computerized protocol. The tip was moved alternately in rostral and caudal directions with displacement in a progressively increasing manner (4-1280 μm) at a speed of 300 μm/s (Figure 3-figure supplement 2A and C). The holding time was set for 2 s between the rostral and caudal movement of the probe tip.

Since all the Aδ-LTMRs were spontaneously silent at the beginning of the experiment without any intentional stimuli, the mechanical response threshold was defined as the displacement that induced the first discharge during the stimulation protocol. If a fiber showed no action potentials during the course of the protocol, even though it exhibited firing as a result of manual touch with a blunt glass rod, then the mechanical threshold was defined to be 1,280 μm. The magnitude of the touch response was represented by the number of spikes evoked during the rostral or caudal movement of the probe (i.e. ramp phases), but not during holding phases of 2 s without probe movement.

### Cold and Heat Stimulation

We applied ramp-shaped thermal stimuli to the RF using a feedback-controlled Peltier device (intercross-2000N, Intercross, Co. Ltd., Japan) with a small probe (diameter: 1 mm). From a baseline temperature of 32°C, the RF was gradually cooled down to 8°C over 40 s or heated up to 50°C over 30 s at a constant rate of 0.6°C/s. Original temperature traces are shown in Figure 3-figure supplement 2B.

### Preparation of Extracellular Matrix Proteins

FLAG-tagged mouse full-length EGFL6 was purified as described below. 293F cells were transfected with the *Egfl6-flag* expression vector and cultured for 4 days according to the manufacturer’s instruction (Invitrogen). The supernatant was collected and incubated with anti-FLAG M2 affinity gel (Sigma) overnight at 4°C. The FLAG affinity gel was collected in an empty Econo-Column (Bio-Rad) and washed with PBS. Bound protein was eluted with 100 μg/ml FLAG peptide in PBS. The eluted protein solution was dialyzed with PBS. Protein concentration was measured with a Pierce BCA Protein Assay Kit using bovine serum albumin (BSA) as a control. The expression vector of RGE-mutant *Egfl6* was generated using the Toyobo KOD-Plus-Mutagenesis kit (Toyobo). FLAG-tagged mouse full-length nephronectin was purified as described previously (Fujiwara et al., 2011). Human plasma fibronectin (Wako) and laminin-511-E8 (Nippi) were purchased from the companies indicated.

### Solid-Phase Cell Adhesion Assays

Solid-phase cell adhesion assays were performed as described previously (Fujiwara et al., 2011) with minor modifications. Briefly, 96-well cell culture plates were coated with purified ECM proteins and blocked with 1% heat-denatured BSA. Human primary and cultured cell lines were suspended in serum-free DMEM, plated on the coated plates, and incubated for 30 min in a CO_2_ incubator at 37°C. Attached cells were fixed and stained with 0.5% crystal violet in 20% methanol for 15 min. The cell-bound crystal violet was extracted with 1% SDS solution and the absorbance was measured at 595 nm. To inhibit cell adhesion activity of different integrins in cell adhesion assays, integrin function blocking antibodies were added to cell suspension before seeding to the coated wells. The cell suspension with the antibodies was incubated for 20 min and then plated on the coated dishes. The function-blocking integrin antibodies used in this study were listed below: integrin α2 (P1E6), α3 (P1B5), α5 (P1D6), α6 (GoH3), αv (L230), and β1 (P5D2). P1E6, P1B5, P1D6, and P5D2 were purchased from the Developmental Studies Hybridoma Bank of the University of Iowa. GoH3 and L230 were purchased from BioLegend and ATCC, respectively. Cell lines used in the assays were human primary keratinocytes, HEK293, T98G, KG-1-C, SF-TY, NTI-4, HT-1080, K562 and K562-a8.

### Hair plucking assays

Dorsal hairs of 20-day-old anesthetised BALB/c mice were dyed using a neon orange hair colour cream for 30 min. Mice were maintained till the tip of new growing undyed hairs could be observed at the skin surface under a dissection microscope (P32-35). The dyed long club hairs, but not short emerging undyed new hairs, were plucked using tweezers under anaesthesia in an area of 10 x 15 mm. Three days after club hair plucking (P36-38), dorsal skin tissue samples containing both plucked and unplucked hair follicles were collected and subjected to whole-mount immunostaining. Plucked and unplucked skin areas could be easily distinguished by hair colour: the unplucked area showed dyed orange hair colour and the plucked area showed undyed white hair colour. In skin tissue samples, hair shafts could be identified by their autofluorescence. This allowed us to distinguish plucked and unplucked hair follicles within tissue samples under a confocal microscope. Zigzag hair follicles were selected for the analysis. Hair follicle types were distinguished by the following criteria under light-field illumination of 150 μm-thick tissue sections. Zigzag hair follicles are the most abundant follicle type in adult mouse dorsal skin (~81%) (Driskell, Giangreco, Jensen, Mulder, & Watt, 2009). Growing new zigzag hairs in plucked and unplucked hair follicles have one row of medulla cells and the smallest diameter (9.58 ± 0.90 μm, n = 32 hair follicles) at the bulge region in comparison to other hair types. Awl and auchene hair follicles together make up ~17% of the dorsal skin hair follicles. Growing awl/auchene hairs in plucked and unplucked hair follicles have more than three rows of medulla cells and have a large hair shaft diameter (15.7 ± 1.95 μm, n = 27 hair follicles) at the bulge region. Guard hair follicles are rare (~1% of the adult dorsal skin hair follicles) and can be clearly identified by their prominent large follicle diameter.

### Statistical analysis

Statistical parameters including the numbers of samples and replicates, types of statistical analysis and statistical significance are indicated in the Results, Figures and Figure Legends. p values: *p < 0.05; **p < 0.01; ***p < 0.001.

### Data availability

Fastq files of RNA-seq data have been submitted to NCBI SRA, and these data can be accessed through the BioProject ID: PRJNA342736.

## Supplemental Figures and Tables

**Figure 1-figure supplement 1.**
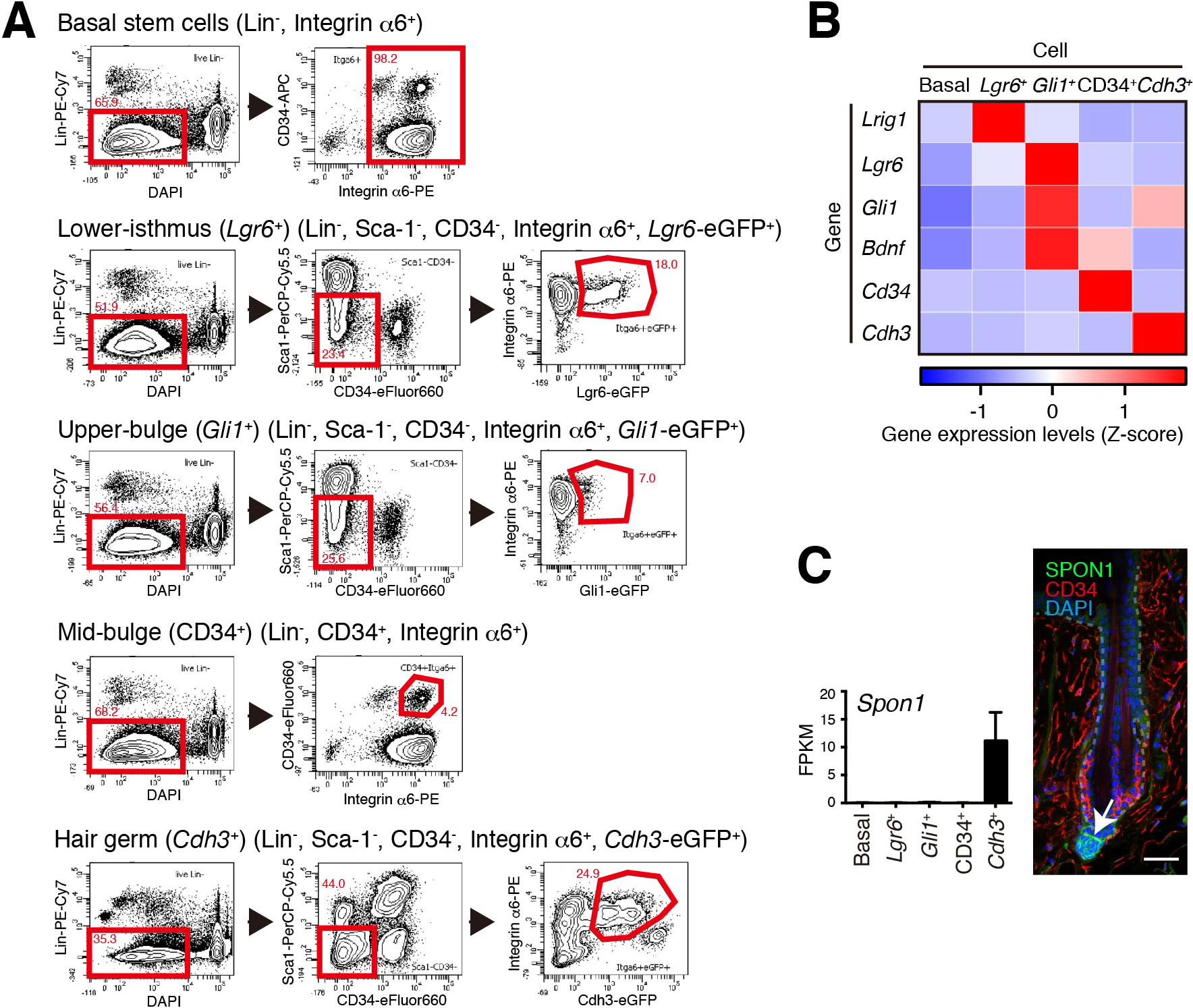
FACS-Based Cell Isolation Procedures for Distinct Epidermal Stem Cell Populations. (A) FACS sorting procedures of all basal epidermal cell populations using wild-type C57BL/6N mice, *Lgr6*^+^ lower isthmus epidermal stem cells using *Lgr6-GFP-ires-CreERT2* mice, *Gli1*^+^ upper-bulge epidermal stem cells using *Gli1-eGFP* mice, CD34^+^ mid-bulge epidermal stem cells using wild-type C57BL/6N mice, *Cdh3*^+^ hair germ epidermal cells using *Cdh3-eGFP* mice. Gates are indicated by red-line boxes and cells in the gates were further analysed in the next plots or sorted. The numbers in the plots represent the percentage of cells in the gates. Lin^−^ indicates lineage-negative cells, which are negative for the markers of haematopoietic and endothelial cells (lineage-positive cells). (B) Z-score heat map representing qRT-PCR analysis of sorted cells with compartment-specific gene primers. See Methods for more detail. Data are mean of 3-4 independently isolated biological replicates. (C) Expression levels of *Spon1* gene in different stem cell pools. Immunostaining pattern of SPON1 protein in 8-week-old telogen dorsal hair follicle was shown. White arrow indicates the restricted localization of SPON1 in dermal papilla and the basement membrane between dermal papilla and hair germ. This restricted expression and deposition of SPON1 corroborates little contamination of hair germ cells into the *Gli1*+ upper-bulge population. FPKM, fragments per kilobase of transcript per million mapped reads. Basal, basal epidermal stem cell pool. Data are mean ± SD, n = 3-4. Scale bar, 10 μm.

**Figure 2-figure supplement 1.**
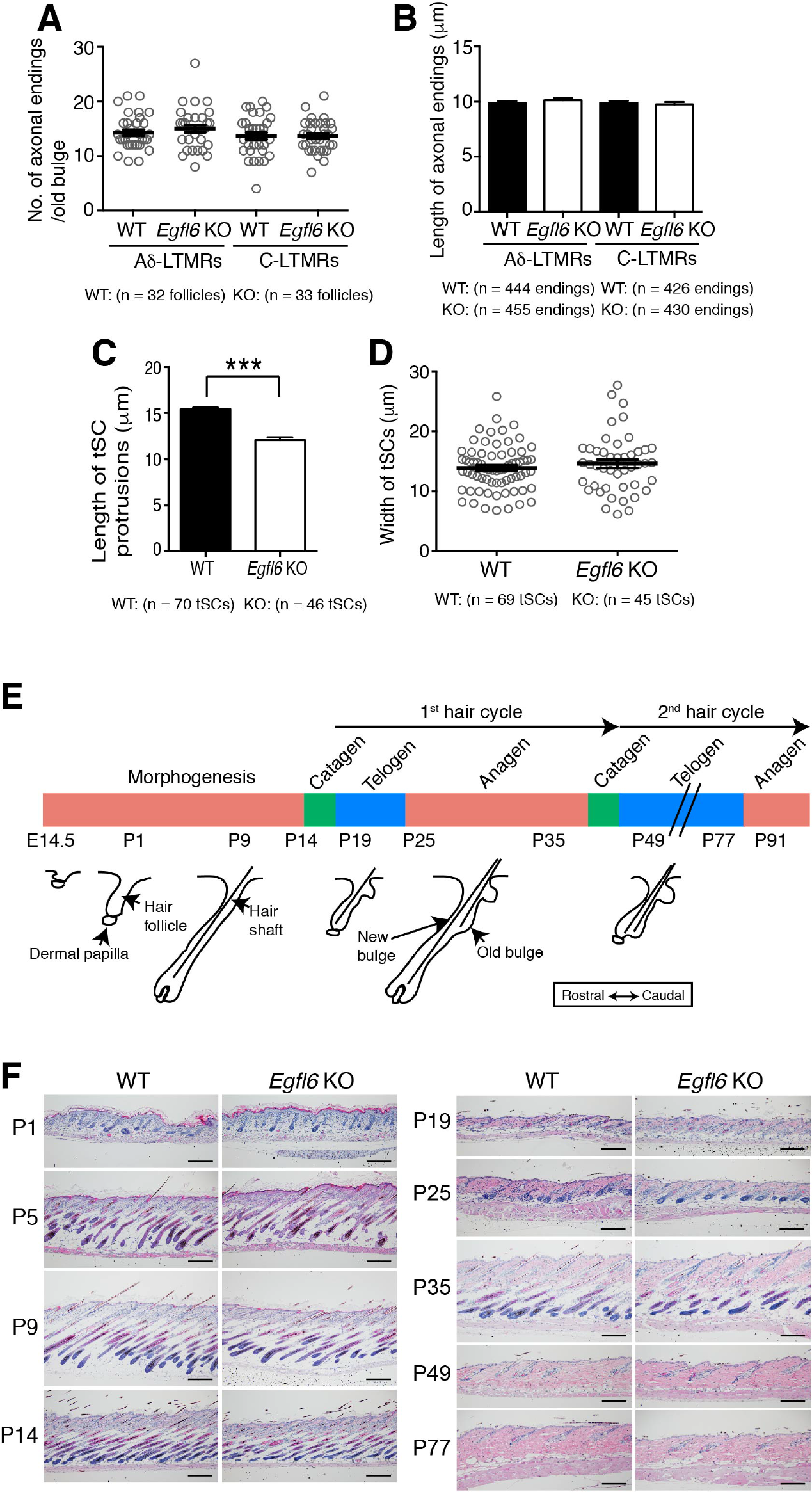
Cell Adhesion to EGFL6 is Mediated by αv Integrins. (A) Coomassie Brilliant Blue (CBB) staining and western blotting of purified EGFL6, RGE-mutant EGFL6, and nephronectin (NPNT) are shown. (B) Heat map display of the results of cell adhesion assays. The number of adhered cells was converted to arbitrary units (A.U.). FN, fibronectin; LN511-E8, laminin-511 E8 fragment. n = 3. Cell lines used are human primary keratinocytes, human embryonic kidney epithelial cell line HEK293, human glioblastoma cell line T98G, Human glioma cell line KG-1-C, human skin fibroblast cell line SF-TY, human embryonic fibroblast cell line NTI-4, human sarcoma cell line HT-1080, human myelogenous leukemia cell line K562 and K562 transfected with human integrin α8 cDNA K562-a8. (C) Cell adhesion assays with T98G and KG-1-C cells plated on EGFL6 or RGE-mutant EGFL6-coated dishes. Data are mean ± SEM, n = 3. (D) Cell adhesion inhibition assays with integrin antibodies. Data are mean ± SEM, n = 3. (**E**) 8-week-old telogen dorsal HFs of wild-type and *Egfl6* knockout mice were stained for EGFL6, integrin αv and nestin. Scale bar, 10 μm. Blue dashed lines indicate the surface of follicle epidermis and white dashed lines indicate cross-section positions. (**F**) Quantification of αv integrin signal intensity in lanceolate complexes. Data are mean ± SD. Statistics (C, D, F): twotailed unpaired *t*-test.

**Figure 3-figure supplement 1.**
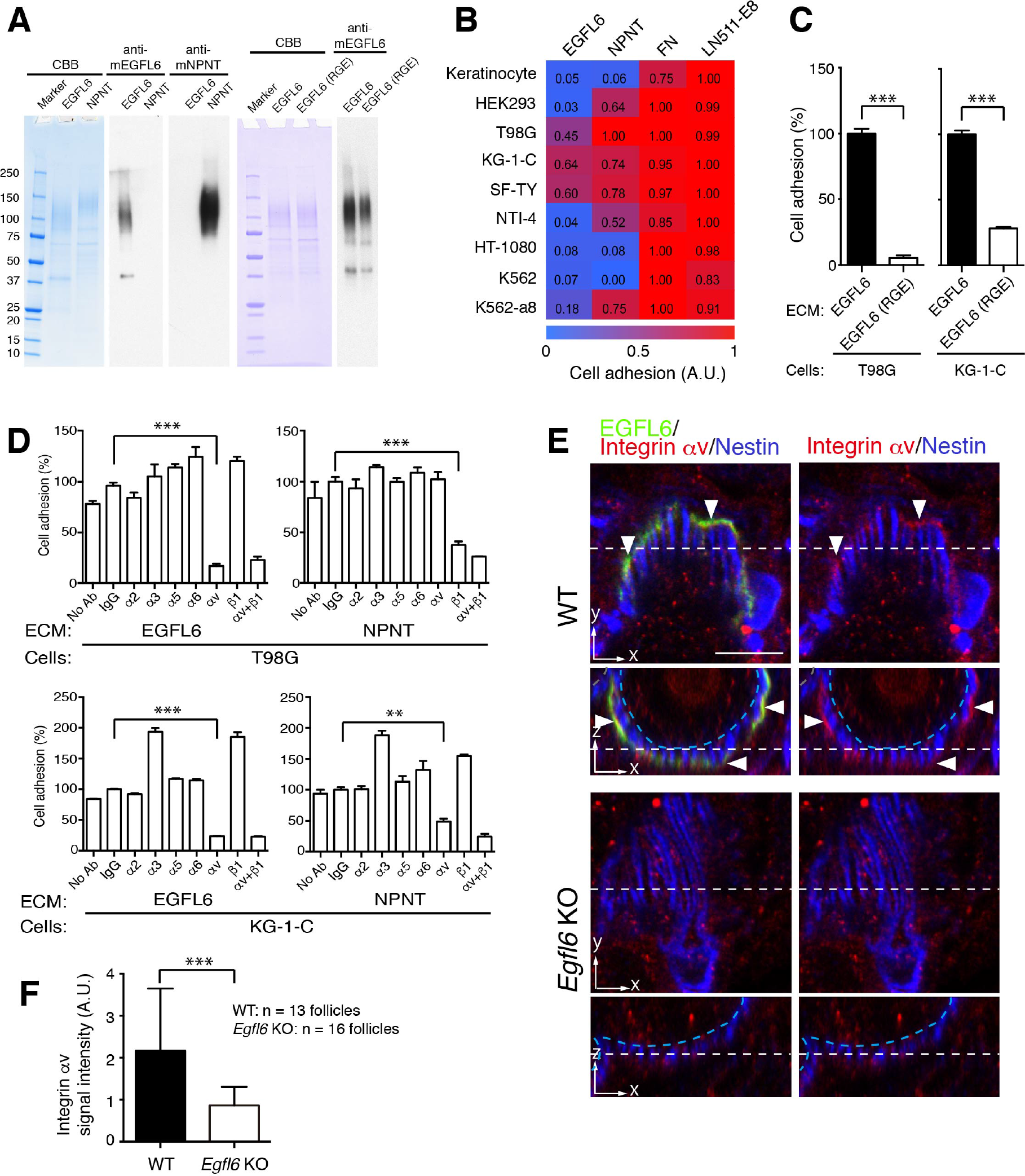
Quantification of Morphological Characteristics of Axonal Endings and tSC Processes and Histological Examination of Skin Tissue Morphology in Wild-type and *Egfl6* Knockout Mice. (A) Number of axonal endings in the old bulge of zigzag hair follicles was counted. Data are mean ± SEM, n = 32 hair follicles for wild-type and n = 33 hair follicles for knockout from 3 mice each. (B) Length of axonal endings in the old bulge of zigzag hair follicles was measured. Data are mean ± SEM, n = 444 axonal endings for wild-type and n = 455 axonal endings for knockout from 3 mice each. (C) Length and width of tSC processes were measured. Data are mean ± SEM, n = 70 tSCs for wild-type and n = 46 tSCs for knockout from 3 mice each. (D) Width of tSC processes were measured. Data are mean ± SEM, n = 69 tSCs for wild-type and n = 45 tSCs for knockout from 3 mice each. (E) Schematic representation of mouse hair follicle morphogenesis and the following regeneration cycles. During the first anagen phase (P25-P35), old and new hair bulge structures form. (F) H&E-stained dorsal skin of wild-type and *Egfl6* knockout mice. Scale bars, 100 μm.

**Figure 3-figure supplement 2.**
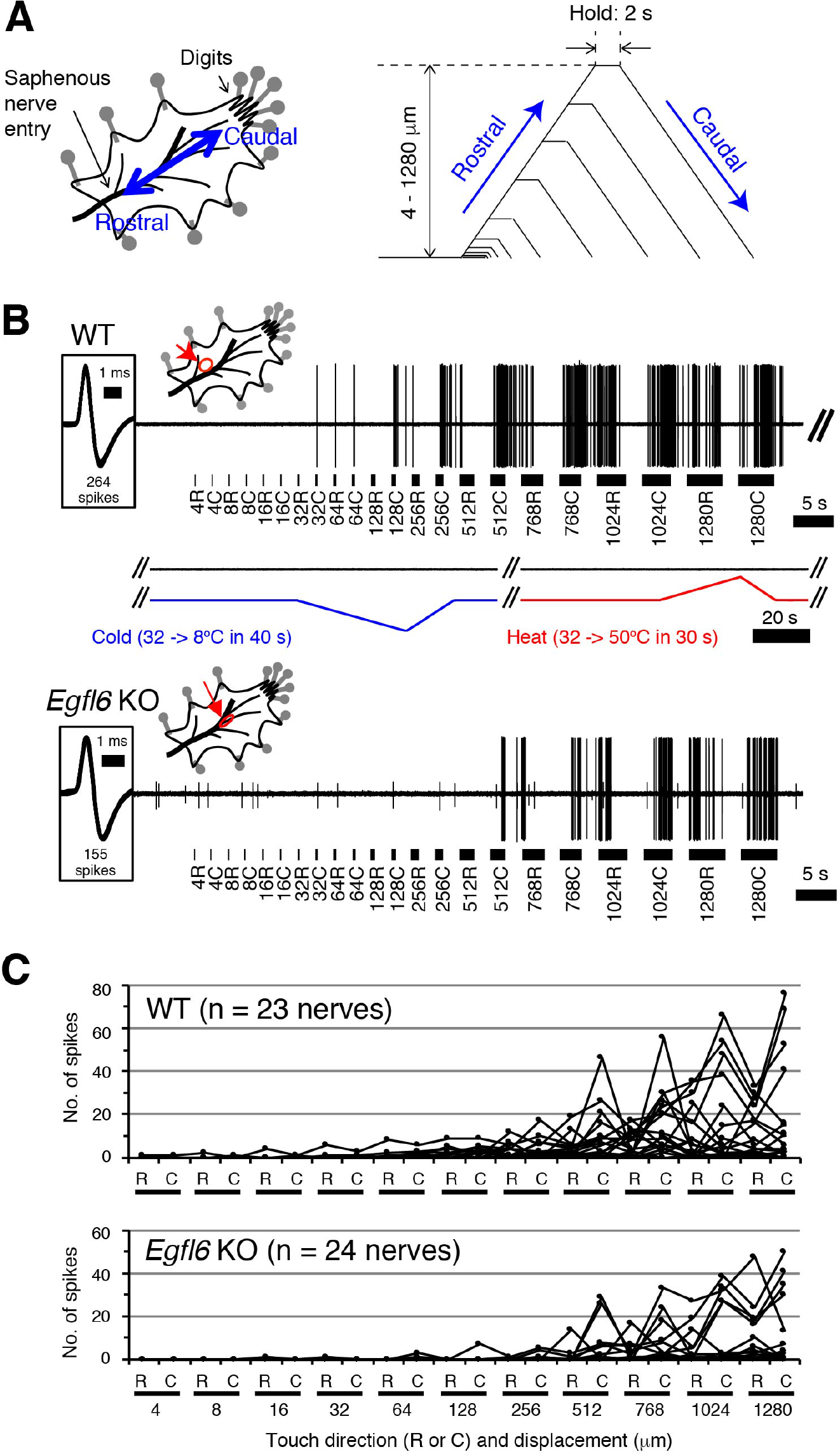
Skin-Nerve Preparation and Recordings of Responses of Aδ-LTMRs. (A) Mouse skin-nerve preparation *ex vivo*, which is oriented with the dermis side up in the test chamber and affixed with pins. A ramp-and-hold touch stimulus protocol is shown. A probe tip was moved alternately in rostral and caudal directions with displacement in a progressively increasing manner (4-1280 μm) at a speed of 300 μm/s. The holding time was set for 2 s between the rostral and caudal movement of the tip. (B) Representative responses in the wild-type (upper panel) and *Egfl6* knockout skin (lower panel). Note the first spike of the wild-type in response to touch stimulus with caudal displacement of 32 μm (i.e. 32C). No firings were observed in response to cold (from 32°C to 8°C in 40 s) or heat (from 32°C to 50°C in 30 s) stimulations. In the knockout, smaller responses to touch stimuli can be observed with the higher threshold displacement of 512 p,m. Spike shapes are shown in insets squared at the left side. All of the spikes generated during a series of stimuli are superimposed (264 spikes in wild-type, and 155 spikes in knockout). The distribution of receptive fields is indicated by an arrow. (C) Individual response patterns in the wild-type (n = 23, upper panel) and the knockout (n = 24, lower panel).

**Figure 4-figure supplement 1.**
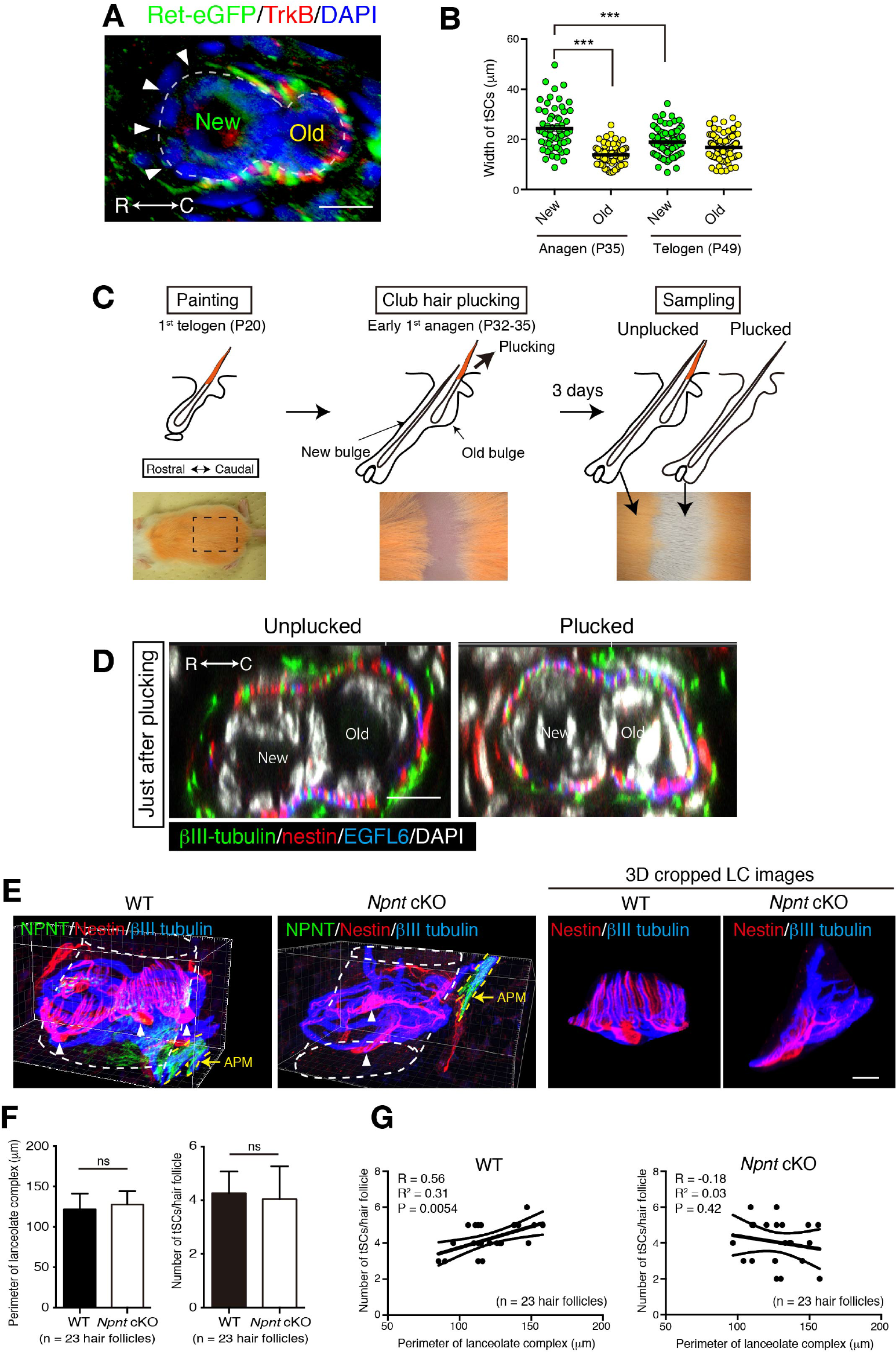
The old bulge formation provides stable epidermal-neuronal interfaces. (A) A cross-sectional view of 7-week-old telogen upper bulge stained for *Ret*-eGFP, TrkB and DAPI. Closed arrowheads indicate a gap of lanceolate endings. (B) The width of tSCs in new and old bulges in the first anagen (P35) and second telogen (P49) follicles were quantified. n = 60-92 tSCs from 3 mice. Individual measurement data are shown as circles. Mean ± SEM are shown as bars, n = 3 mice. Statistical comparisons were performed using two-tailed unpaired t-test. (C) Procedure of club hair plucking experiments. (D) Transverse sectional views of upper-bulge with club-hair unplucked and plucked P32 early anagen hair follicles stained for βIII-tubulin, nestin, EGFL6 and DAPI. Just after plucking, little structural damage to the old bulge epidermis and LCs were observed. (E) 3D reconstituted immunostaining images of upper-bulge region of 7-week-old telogen Awl/auchene hair follicles of wild-type and epidermis-specific conditional nephronectin (*Npnt*) knockout mice (Keratin5-Cre/Npnt fl/fl). Upon epidermal deletion of *Npnt* gene, arrector pili muscles (APM) moved above the bulge (Fujiwara et al., 2011), where LCs were located. Arrowheads indicate the nuclei of tSCs. In the mutant, some hair follicles showed disrupted patterning of longitudinal sensory nerve endings and tSCs processes. This mutant image shows a significantly fewer number of longitudinal lanceolate endings and tSC processes (see also 3D cropped LC images). (F) The perimeter and number of LCs in randomly selected non-guard hair follicles of wild-type and *Npnt* conditional knockout mice were measured. No statistically significant difference was observed. Data are mean ± SD, n = 23 hair follicles from wild-type and mutant mice, respectively. (G) The number of tSCs were plotted against the perimeter of LCs. In the wild-type, the number of tSCs correlated well with the perimeter of LCs, *i.e*. perimeter of hair follicles, while in the *Npnt* knockout mice, distribution of the values of “number of tSC/perimeter of LCs” was scattered and their correlation was lost (see R^2^ and p-value), indicating the anatomical changes of LCs in *Npnt* mutants. n = 23 hair follicles from wild-type and mutant mice, respectively. Scale bars, 10 μm.

**Figure 1- table supplement 1.**
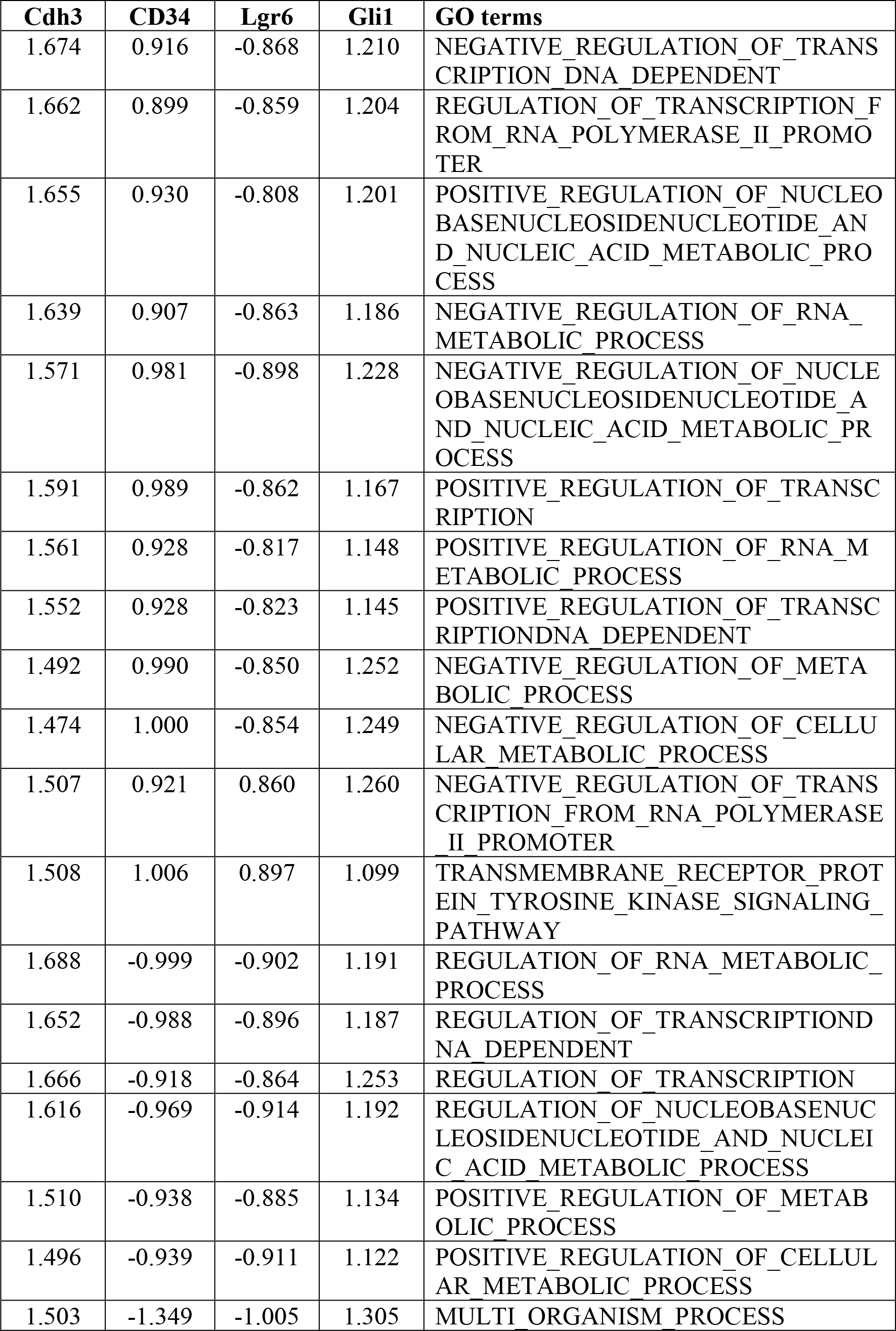
Normalized Enrichment Score (NES) of Gene Set Enrichment Analysis (GSEA)

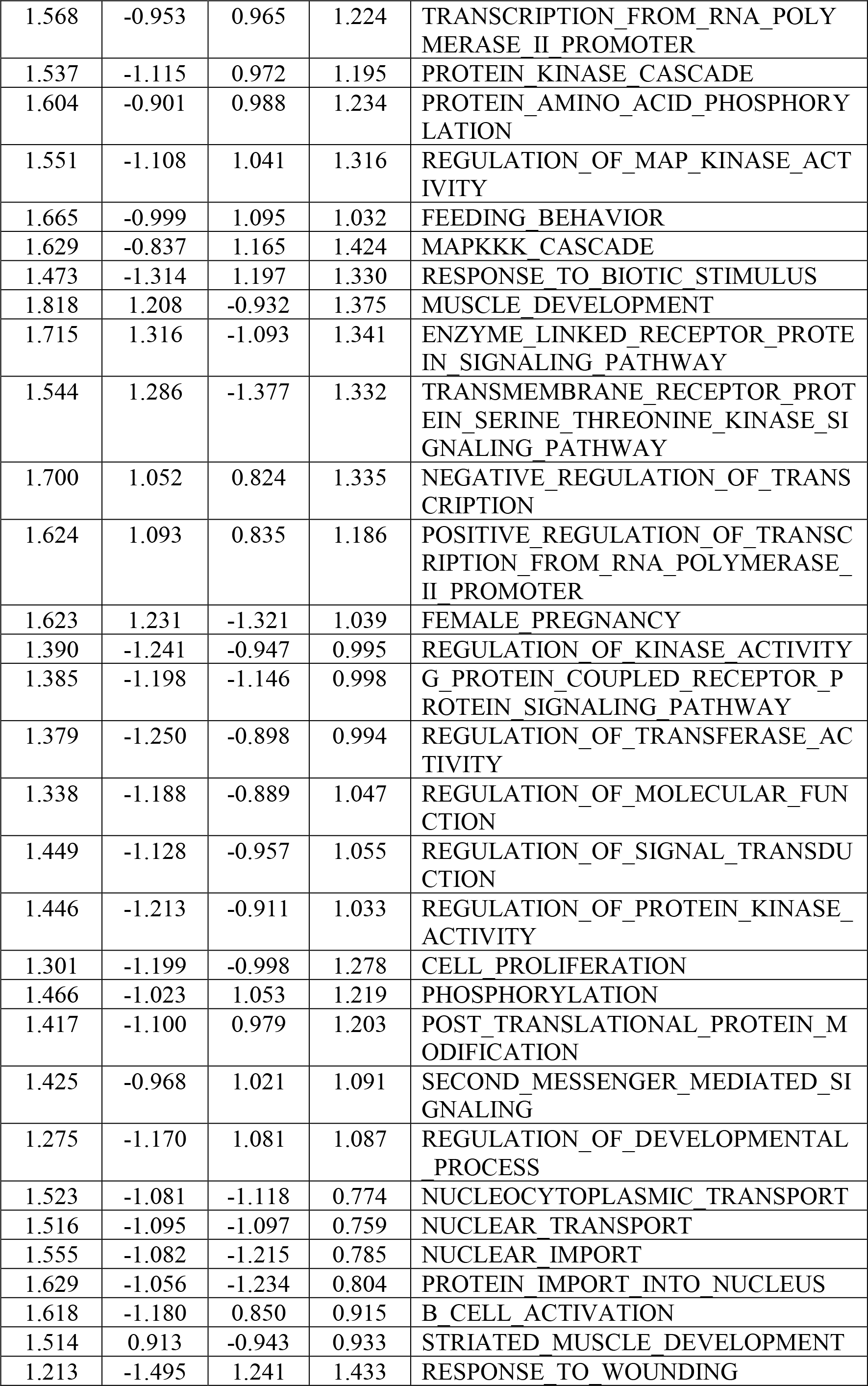

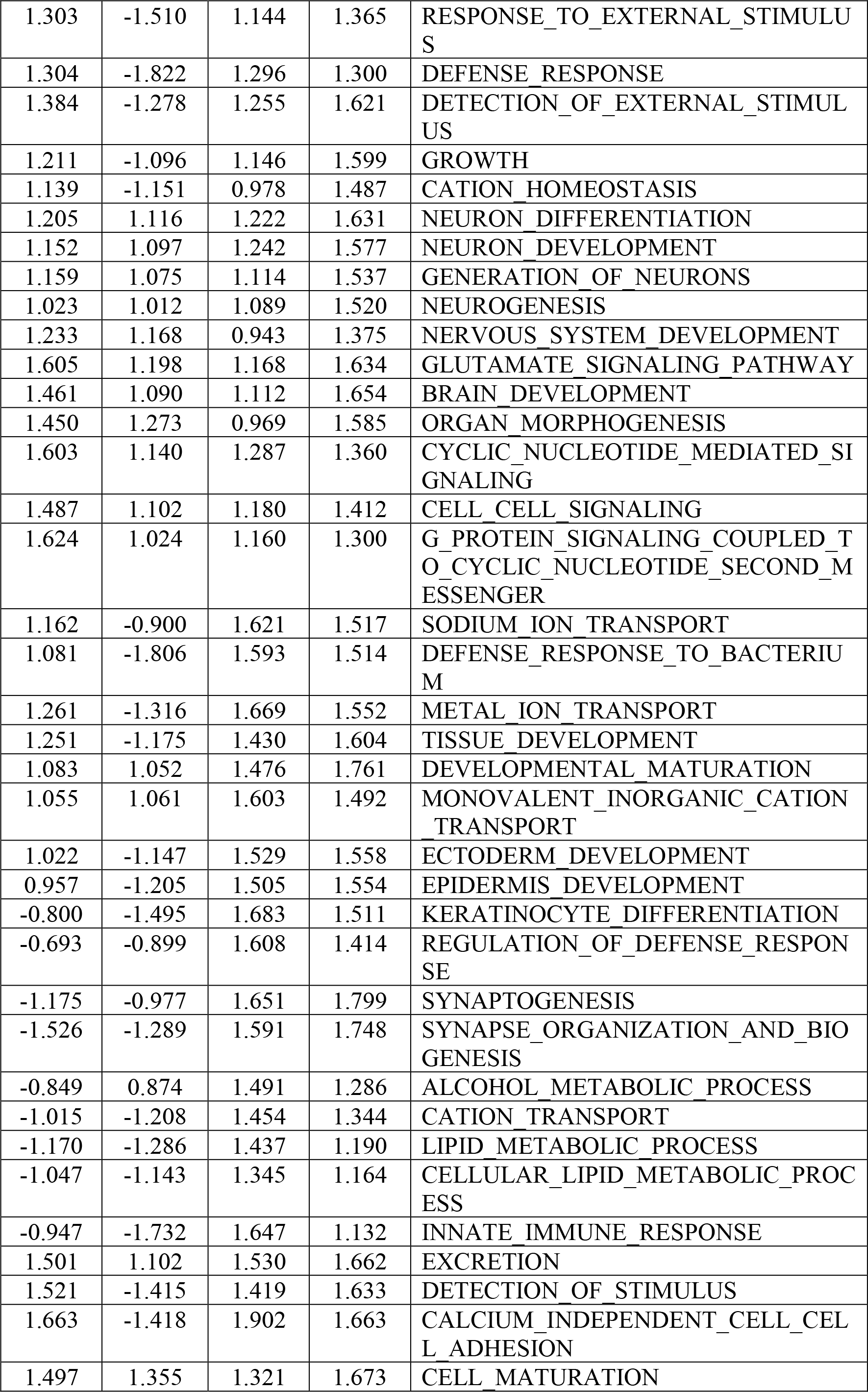

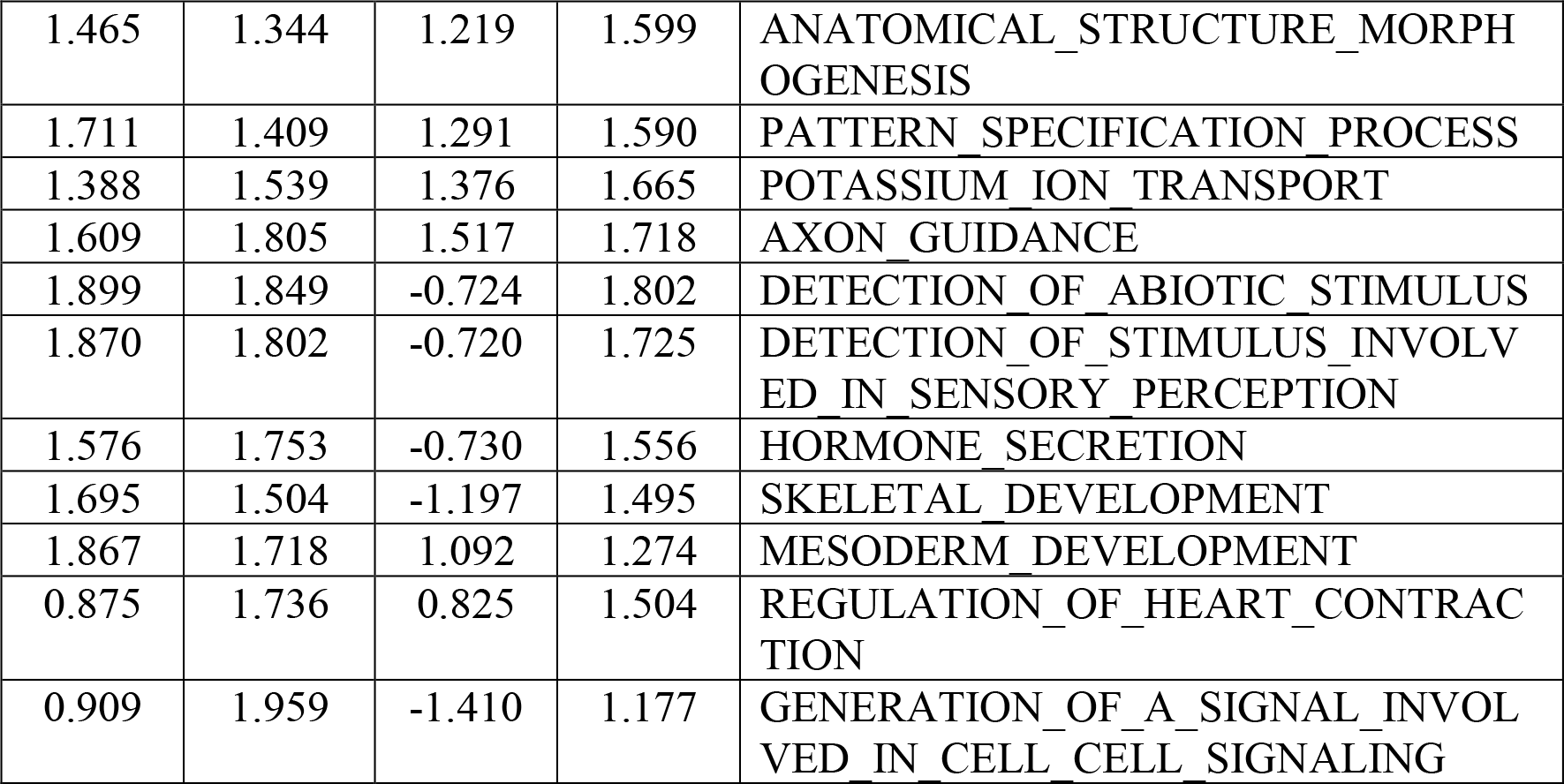

**Figure 2-table supplement 1.** List of ECM proteins screened for upper-bulge localization with immunohistochemical analysis.

| Gene symbol | Protein name (Abbreviation) | Antibody ID | Manufacturer | Upper-bulge localization | Tissue localization |
| --- | --- | --- | --- | --- | --- |
| Aspn | Asporin | ab58741 | abcam | Yes | Upper bulge sensory neuron |
| Crispld1 | Cysteine-rich secretory protein LCCL domain containing 1 (CRISPLD1) | ab123039 | abcam | Yes | Basement membrane zone of upper-bulge |
| Epdr1 | Ependymin related protein 1 (EPDR1) | PA283664 | Cusabio | No | Uncertain |
| Col5a2 | Collagen, type V, alpha 2 (COL5A2) | ab7046 | abcam | Yes | Broadly distributed in the epidermal-dermal basement membrane zone |
| Vwa2 | von Willebrand factor A domain-containing protein 2 (VWA2) | ab111164 | abcam | No | Follicle-arrector pili muscle junction |
| Fam101b | Refilin B | orb183474 | biorbyt | No | Uncertain |
| Ltbp1 | Latent TGF-beta-binding protein 1 (LTBP1) | PA807018 | Cusabio | No | Uncertain |
| Postn | Periostin | ab14041 | abcam | Yes | Basement membrane zone of upper-and mid-bulge |
| Spon2 | Spondin-2 | PA006509 | Cusabio | No | Uncertain |
| Egfl6 | Epidermal growth factor-like protein 6 (EGFL6) | CUK1203 | Fujiwara lab (in house) | Yes | Upper-bulge collar matrix |
| Igfbp5 | Insulin-like growth factor binding protein 5 (IGFBP5) | AF578 | R&D | Yes | Upper bulge sensory neuron |
| Col8a2 | Collagen, type VIII, alpha 2 (COL8A2) | 34099 | USBiological | No | Uncertain |
| Igfbp7 | Insulin-like growth factor binding protein 7 (IGFBP7) | ab129302 | abcam | No | Uncertain |
| Adamtsl4 | ADAMTS-like 4 (ADAMTSL4) |  |  | Not tested |  |
| Col4a3 | Collagen, type IV, alpha 3 (COL4A3) | H31 | Shigei Med. Res. Inst. | Yes | Basement membrane zone of upper-bulge |
| Col4a4 | Collagen, type IV, alpha 4 (COL4A4) | RH42 | Shigei Med. Res. Inst. | Yes | Basement membrane zone of upper-bulge |

**Figure 3-table supplement 1.** General characteristics of Aδ-LTMRs.

| Outcomes | WT<br>(n=23 nerves) | *Egfl6* KO<br>(n=24 nerves) | *p* value |
| --- | --- | --- | --- |
| Activation threshold (V) | 3.0 (2.0-4.0) | 3.0 (2.0-6.0) | *p* = 0.778 |
| Conduction velocity (m/s) | 6.6 (5.1-7.7) | 7.7 (5.0-8.6) | *p* = 0.115 |
| Spontaneous activity (imp/s) | 0 (0-0) | 0 (0-0) | *p* > 0.999 |
| Receptive field size (mm<sup>2</sup>) | 6.5 (4.7-13.2) | 4.8 (2.8-7.3) | *p* = 0.059 |

